# Innate immune signaling and sex differences contribute to neurocognitive impairment, neuroinflammation, and mitochondrial rewiring in a mouse model of Gulf War illness

**DOI:** 10.1101/2020.08.28.271833

**Authors:** Joshua D. Bryant, Maheedhar Kodali, Bing Shuai, Saeed S. Menissy, Paige J. Graves, Ashok K. Shetty, Laura Ciaccia West, A. Phillip West

## Abstract

Gulf War Illness (GWI) is a chronic, multi-symptom disorder affecting approximately 30 percent of the nearly 700,000 veterans of the 1991 Persian Gulf War. Recent studies have revealed that GWI-related chemical (GWIC) exposure promotes immune activation and metabolic rewiring, which correlate with neurocognitive impairments and other symptoms of GWI. However, the molecular mechanisms and signaling pathways linking GWIC to inflammation, metabolic alterations, and neurological symptoms remain unclear. Mitochondrial dysfunction has been documented in veterans with GWI and rodent models, and because mitochondria are key immune regulators, we hypothesized that alterations to mitochondria-immune crosstalk could contribute to the development of GWI-related symptoms. Here we show that acute exposure of murine macrophages to GWIC alters mitochondrial respiration and potentiates innate immune signaling and inflammatory cytokine secretion. Using an established mouse model of GWI, we report that neurobehavioral changes, neuroinflammation, and mitochondrial protein rewiring are attenuated in mice lacking the cyclic GMP-AMP synthase (cGAS)-Stimulator of Interferon Genes (STING) and NOD-, LRR- or pyrin domain-containing protein 3 (NLRP3) innate immune pathways. Finally, we report sex differences in response to GWIC, with female mice showing more pronounced cognitive impairment, neuroinflammation, and mitochondrial protein alterations in the brain compared to male mice. Our results provide novel information on sex differences in this model and suggest that STING and NLRP3 are key mediators of the cognitive impairment, inflammation, and mitochondrial dysfunction observed in GWI.

## 1. Introduction

Gulf War illness (GWI) is a chronic, multi-symptom disorder affecting approximately 25 to 32 percent of the nearly 700,000 veterans of the 1991 Persian Gulf War (PGW-1). Predominantly characterized by brain-related problems including learning and memory deficits, depression, and anxiety, GWI has been linked to synergistic exposure to chemicals including the nerve gas prophylactic pyridostigmine bromide (PB), a reversible acetylcholinesterase inhibitor, and permethrin (Per), an insecticide^1,2^.

Research over the past few decades has uncovered mechanisms by which exposure to GWI-related chemicals (GWIC) cause GWI pathology and symptoms. Normal function of the brain and nervous system requires significant cellular energy, which is generated within cells by mitochondria. GWIC exposure enhances reactive oxygen species (ROS) production and oxidative stress^3^, which may contribute to a vicious cycle of mitochondrial dysfunction (MD) by damaging mitochondrial DNA (mtDNA) and oxidative phosphorylation (OXPHOS) proteins to further augment ROS and trigger progressively increasing mitochondrial stress. Multiple studies have documented MD in both human GWI patients^4,5,6^ and in murine models of GWI^3,7,8,9^. Moreover, clinical manifestations of GWI resemble those identified in Chronic Fatigue Syndrome (CFS), which has been linked with MD^5,10^.

Immune system alterations have also been associated with GWI pathology. Studies of GWI patients have revealed immune changes, including increases in inflammatory cell types^11–13^, heightened production of proinflammatory cytokines^11,14–16^, and transcriptional profiles that resemble those observed in autoimmune disorders^17^. Parallel immune alterations have also been observed in rodent models of GWI, with increases in proinflammatory cytokines seen in the hippocampi of GWIC-exposed rats^3^ and in the brains and serum of GWIC-exposed mice^9,18^. However, the role of the innate immune system in GWI pathology, and specifically the identification of signaling cascades responsible for proinflammatory cytokine induction in animal models and Veterans with GWI, remains uncharacterized.

Several GWI studies have demonstrated MD and inflammation contemporaneously, suggesting an interaction between these two disease mechanisms^19^. Indeed, in addition to their well-appreciated roles in cellular metabolism and apoptosis, mitochondria are key immune regulators, impacting many aspects of signaling and effector responses in both the innate and adaptive immune system^20^. MD and mtDNA damage can trigger a wide range of inflammatory responses that have been implicated in immune-mediated pathology^21^.

Based on evidence of MD and inflammation in GWI, and on the documented link between the two, we hypothesized that exposure to GWIC might alter mitochondria-innate immune crosstalk, contributing to inflammatory responses that exacerbate neuropathology and cognitive symptoms of GWI. To test this hypothesis, we employed an established mouse model of GWI in which mice are exposed to the GWIC by intraperitoneal injection for a period of 10 days, developing cognitive impairment and neuroinflammation beginning at 5 months post-exposure^22–24^. In our study, we employed a longitudinal analysis regimen, performing neurocognitive tests at both 5 months and 12 months after acute exposure to GWIC, followed by analysis of mitochondrial protein expression and neuroinflammation in the brain at the terminal timepoint. We used both male and female mice in these experiments to evaluate sex-dependent effects of GWIC, as there is evidence that GWI can be more severe in women^25^. To examine the acute effects of GWIC exposure on metabolism and cell-intrinsic immune responses, we performed a series of *in vitro* experiments using bone marrow-derived macrophages. And finally, we applied this GWIC-exposure model to C57BL/6 genetic knockouts lacking key innate immune sensors that interface with mitochondria and cellular metabolism to evaluate the contributions of these inflammatory signaling pathways to development of GWI pathology. Our results provide important new information on sex differences in GWI-related pathology and reveal that innate immune signaling contributes to neurocognitive impairment, neuroinflammation, and mitochondrial rewiring in a preclinical model of GWI.

## 2. Materials and Methods

### 2.1 Animals, study design, and Gulf War chemical agents

All C57BL/6J wild-type (WT) mice were purchased directly from Jackson Laboratories and allowed to acclimate in the animal facility for 2-4 weeks, or bred in-house. Breeding pairs of NLR Family Pyrin Domain Containing 3 null (NLRP3^−/−^) (JAX strain 021302) and Stimulator of Interferon Genes null (STING^gt/gt^) (JAX strain 017537) mice were purchased from Jackson Laboratories and bred to generate sufficient mice for all experiments. Mice were 8-10 weeks of age upon exposure to GWIC.

Briefly, mice were randomly assigned to treatment cohorts: DMSO (vehicle control; 15 mice per sex for WT mice, 6 mice per sex for NLRP3^−/−^ and STING^gt/gt^ null mice) or GWIC (15 mice per sex for WT, NLPR3 null, and STING^gt/gt^ mice). Mice then underwent an exposure regimen of daily peritoneal injections of 50 microliters of DMSO alone for DMSO cohorts or 50 microliters of GWIC (0.7 mg/kg of PB and 200 mg/kg of Per) dissolved in DMSO for GWIC cohorts. Five months after exposure, 6-10 randomly selected mice from DMSO-treated WT and GWIC-treated WT, NLRP3^−/−^, and STING^gt/gt^ cohorts and all DMSO-treated NLRP3^−/−^ and STING^gt/gt^ mice were subjected to neurobehavioral tests as described below. Twelve months after exposure, mice that were behaviorally tested at five months were retested, and then all mice were sacrificed for tissue processing or immunohistochemical analysis.

The Animal Care and Use Committee (IACUC) of the Texas A&M University College of Medicine, which ensures compliance with all applicate federal and state regulations for the purchase, transportation, housing, and ethical use of animals for research, has approved all experiments performed in this study. The euthanasia methods employed in the study were consistent with the recommendations of the American Veterinary Medical Association (AVMA).

Pyridostigmine bromide (99.9% purity) was purchased from Selleckchem (catalog number HY-B0887) and permethrin (>98% purity) was purchased from MedChemExpress (catalog number S1608).

### 2.2. Behavioral tests for analyses of cognitive and mood function

#### 2.2.1. Object location test (OLT)

The object location test (OLT) involved three sequential trials (Figure 1A top panel): in the first trial, termed the habituation trial, the mouse was placed at the corner of an empty open field apparatus and allowed to freely explore for five minutes; in the second trial (“sample trial”), which took place twenty minutes after the first trial, the mouse was placed in the corner of the same open field apparatus, which now contained two identical objects placed on opposite sides of the box, and allowed to free explore for five minutes; and in the third, “test” trial, after an inter-trial interval of 20 minutes (5 months post-exposure) or 2 hours (12 months post-exposure), the mouse was placed in the corner of the open field apparatus, with one of the objects remaining in its original location and the other object moved to a novel location. The apparatus was cleaned with 70% ethanol and allowed to air-dry prior to the commencement of each trial. The movement of the mouse in the sample and test trials was continuously video-tracked using the Noldus Ethovision XT program; the nose-point of the mouse coming within 2 cm of an object was recorded by the software as object exploration. Data on time spent exploring both objects, total distance traveled, and velocity of movement during these trials were collected. Percent time spent with the object in a novel location was calculated as total time exploring this object divided by total time exploring both objects. Data in which mice spent less than 10 seconds total exploring both objects in either the sample or test trial were excluded.

**Figure 1.**
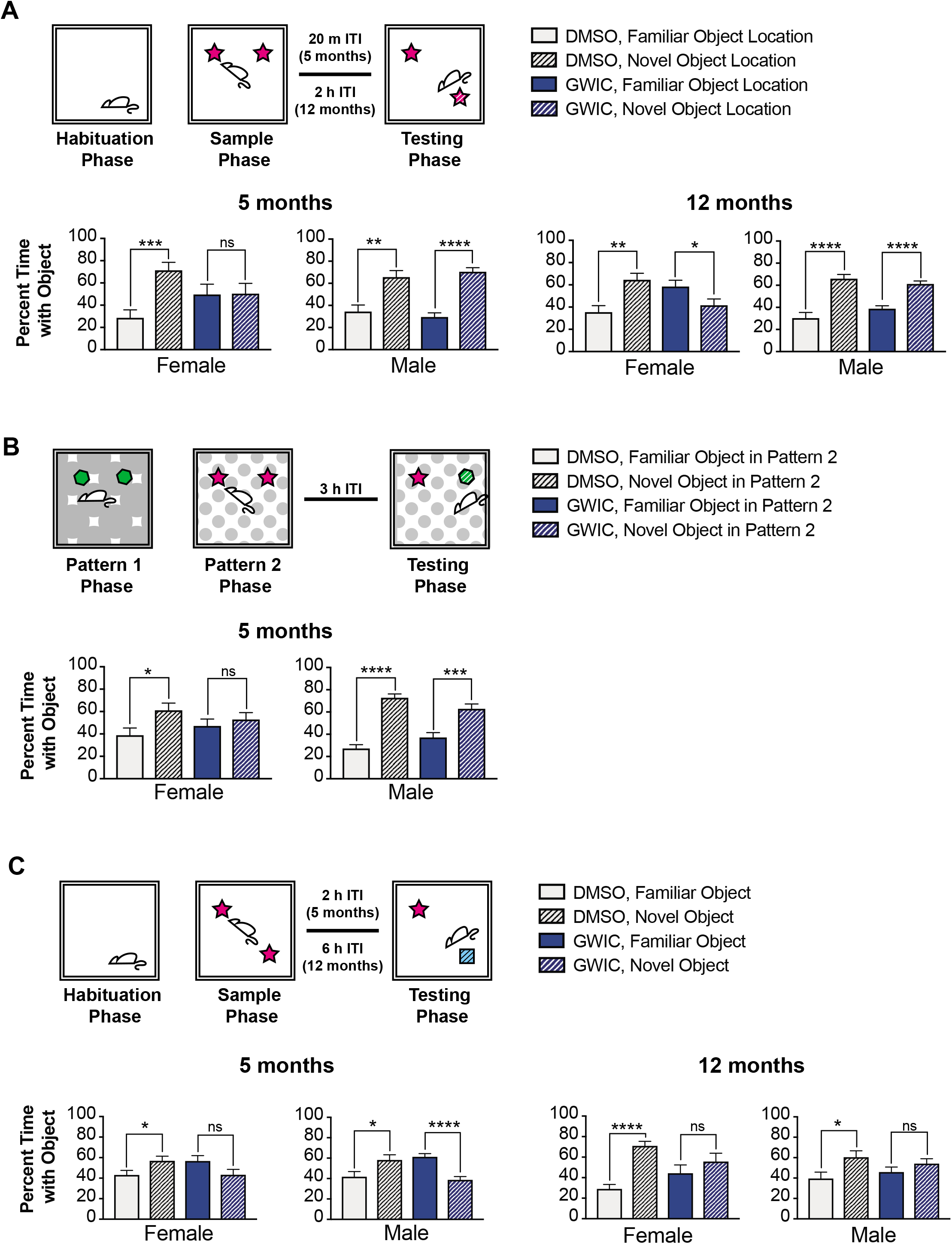
Exposure of C57BL/6 wild-type mice to GWIC results in sex-dependent impairment in spatial memory, pattern separation, and novel object reocognition. (A) The diagram in the top panel depicts the sequential phases used in the object location test (OLT): the habituation phase, the sample phase, and the testing phase. Each phase lasted five minutes, and the inter-trial interval (ITI) between the sample and testing phase was 20 minutes (5 months post-exposure) or 2 hours (12 months post-exposure). The bar graphs in the bottom panel compare the percentages of total exploration time spent with the object in the familiar location versus time spent with the object in the novel location for both the OLT conducted 5 months and 12 months post-exposure, for DMSO- and GWIC-exposed female and male mice. (B) The diagram in the top panel depicts the sequential phases used in the pattern separation test (PST): the Pattern 1 phase, the Pattern 2 phase, and the testing phase, in which one of the objects from Pattern 1 was placed into Pattern 2. Each phase lasted five minutes, and the inter-trial interval (ITI) between the sample and testing phase was 3 hours. The bar graphs in the bottom panel compare the percentages of total exploration time spent with the familiar object in Pattern 2 versus time spent with the novel object in Pattern 2 for a PST conducted 5 months post-exposure, for DMSO- and GWIC-exposed female and male mice. (C) The diagram in the top panel depicts the sequential phases used in the novel object recognition test (NORT): the habituation phase, the sample phase, and the testing phase. Each phase lasted five minutes, and the inter-trial interval (ITI) between the sample and testing phase was 2 hours (5 months post-exposure) or 6 hours (12 months post-exposure). The bar graphs in the bottom panel compare the percentages of total exploration time spent with the familiar object versus time spent with the novel object both the NORT conducted 5 months and 12 months post-exposure, for DMSO- and GWIC-exposed female and male mice.). Error bars represent the mean of biological replicates +/- SEM (n=9-10). Indicated p-values were calculated two-tailed, unpaired, Student’s t-test. *p < 0.05; **p < 0.01; ***p < 0.001; ****p < 0.0001; ns, not significant.

#### 2.2.2. Pattern separation test (PST)

This test also involved three sequential trials, with a few key changes (Figure 1B, top panel). First, the mouse spent five minutes exploring an empty open field that had a graphic black and white pattern (P1) on the floor of the field and two identical objects. Next, the mouse spent five minutes exploring an empty open field that had a different graphic black and white pattern (P2) on the floor of the field and two different identical objects. Following an inter-trial interval of 3 hours, the mouse spent five minutes in a test trial, in which the P2 pattern was on the floor with one object from the P2 phase (familiar) and one object from the P1 phase (novel). The behavior of the mice was video-tracked as described for the OLT, with data collected on the time exploring the novel and familiar objects, distance traveled, and velocity of movement. Percent time spent with the novel object in P2 was calculated as total time exploring the novel object divided by total time exploring both objects. Data in which mice spent less than 10 seconds total exploring both objects in any trial were excluded.

#### 2.2.3. Novel object recognition test (NORT)

This test involved three sequential trials (Figure 1C, top panel): an initial habituation trial as described for the OLT, a second sample trial in which the mouse explored the open field containing two identical objects, and, following an inter-trial interval of 2 hours (5 months post-exposure) or 5 hours (12 months post-exposure), a test trial in which one of the original objects was replaced with a novel object placed in the same location The behavior of the mice was video-tracked as described for the OLT, with data collected on the time exploring the novel and familiar objects, distance traveled, and velocity of movement. Percent time spent with the novel object was calculated as total time exploring the novel object divided by total time exploring both objects. Data in which mice spent less than 10 seconds total exploring both objects in either the sample or test trial were excluded.

#### 2.2.4. Elevated plus maze (EPMT)

The EPMT uses a grey colored maze consisting of a 60 cm elevated plus-shaped apparatus with four arms, each 35 cm in length and 5 cm in width. Two of the arms are surrounded with walls on three sides of 15 cm in height (“closed arms”) and two of the arms are open (“open arms”) (Supplemental Figure 2A). Mice were gently placed in the center of the platform, facing a closed arm, and allowed to freely explore the maze for one five minute interval. The movement of the mouse in the sample and test trials were continuously video-tracked using the Noldus Ethovision XT program, with data collected on: the time that the mouse center-point was observed in open arms, closed arms, and the center of the maze; number of visits to open arms, closed arms, and the center of the maze; distance traveled; and velocity of movement.

### 2.3. Immunohistochemical methods

Twelve months after exposure to GWIC, mice were perfused with 4% paraformaldehyde. Brains were postfixed for 18 hours in 4% paraformaldehyde and cryoprotected in 30% sucrose. A cryostat was then used to cut thirty-micrometer thick coronal sections through the entire brain. The sections were collected serially in 24-well plates containing phosphate buffer; sections through the cerebral cortex at hippocampal levels were chosen and processed for immunohistochemical quantification of glial fibrillary acidic protein (GFAP)-positive astrocytes and ionized calcium-binding adaptor molecular 1 (IBA-1)-positive microglia. Briefly, sections were incubated in 20% methanol and 3% hydrogen peroxide in phosphate buffered saline (PBS) for 20 minutes, rinsed three times in PBS, then incubated for 30 minutes in PBS containing 0.1% Triton-X 100 and 10% serum (Sigma). Following this, sections were incubated in primary antibody (rabbit anti-GFAP antibody or goat anti-IBA-1 antibody, both from Abcam) for 18-48 hours, then biotinylated secondary antibody (biotinylated horse anti-goat IgG or biotinylated goat anti-rabbit IgG from Vector Laboratories) for 60 minutes, then an avidin/biotin complex reagent (Vectastain Elite ABC HRP Kit) for 60 minutes, with a triple wash in PBS between each incubation step. The peroxidase reaction was developed using Vector SG HRP substrate (Vector Laboratories), then sections were mounted on gelatin-coated slides, dehydrated, cleared, and coverslipped.

### 2.4. Analysis of hippocampal inflammation

Astrocyte hypertrophy was evaluated using ImageJ to calculate area fraction of GFAP+ structures in three regions (CA1, CA3, and dentate gyrus) of the hippocampus in six mice per genotype per sex, for DMSO- and GWIC-exposed cohorts. Multiple sections were imaged per mouse and averaged; image collection and ImageJ analysis were blinded to treatment group. Microglia activation was evaluated by performing morphological analysis. Microglia were identified as IBA-1 structures, while activated microglia were identified as IBA-1+ structures with swollen, ramified cell bodies and short, thick processes^26^. Total microglia and activated microglia were counted for three regions (CA1, CA3, and dentate gyrus, DG) of the hippocampus. Percentages of activated microglia among total microglia were computed and compared. Multiple sections were imaged per mouse and averaged; image collection and morphologic analysis was blinded to treatment group.

### 2.5 Tissue processing and expression analysis

For collection of tissues, animals were sacrificed and brains were quickly removed. The entire hippocampus from both hemispheres and cerebral cortex tissues were micro-dissected from each brain, then snap frozen on dry ice. Cells and tissues were lysed in 1% NP-40 lysis buffer supplemented with protease inhibitor and then centrifuged at 12,000 x g, 4°C to obtain cellular lysate. After BCA protein assay (Thermo Fisher Scientific), equal amounts of protein (10–20 μg) were separated on 10-20% SDS–PAGE gradient gels, then transferred onto PVDF membranes. After air drying to return to a hydrophobic state, membranes were incubated in primary antibodies (Table 1) at 4°C overnight in 1X PBS containing 1% casein and 0.02% sodium azide. After incubating with HRP-conjugated secondary antibody at room temperature for 1 hour, membranes were developed with Luminata Crescendo Western HRP Substrate (Millipore). Densitometry was performed in ImageJ and normalized to the GAPDH expression level.

### 2.6 In vitro bone marrow-derived macrophage experiments

L929 cells were obtained from ATCC and maintained in DMEM (D5796) (Sigma) supplemented with 10% FBS (97068-085) (VWR). Bone marrow-derived macrophages (BMDMs) were generated from bone marrow collected from femurs and tibiae isolated from male and female mice. Briefly, legs were dissected from mice and muscle and tissue were removed. Bones were immersed in 70% EtOH then washed with PBS. Bones were crushed with a mortar and pestle and the marrow was collected in DMEM. The marrow was passed through a 70 μm mesh strainer, then blood cells were lysed using ACK lysis buffer. Remaining cells were passed through a 40 μm mesh strainer and plated on Petri dishes in DMEM containing 10% FBS and 30% L929 culture media for 7 days. For all experiments, BMDMs were plated at a density of 6 x 10^5^ cells/mL unless otherwise indicated. After cells were plated and allowed to adhere for 2-3 hours, cells in GWIC treatment group were treated overnight with permethrin (10 μM final) and/or pyridostigmine bromide (PB, 100 ng/mL) as indicated. LPS treatment was done with LPS-B5 Ultrapure (InvivoGen) at a concentration of 200 ng/mL unless otherwise indicated. Transfection of HSV-60bp interferon-stimulatory DNA (ISD) (InvivoGen) into the cytosol of BMDMs was performed using Polyethyleneimine (PEI) (43896) (Alfa Aesar).

### 2.7 ELISA

Primary, secondary, and detection antibodies were purchased from BioLegend. Detached cells and debris were removed from cell culture supernatant by centrifugation prior to the assay. After incubating half-well ELISA plates in capture antibodies at 4°C overnight, plates were blocked with PBS containing 10% FBS at room temperature for 1 hour. Cell culture supernatants were diluted 80-240x (for IL-6) or 2-20x (for TNF-α) in blocking buffer. Standards and cell culture supernatant were added to the ELISA plates and incubated at room temperature for 2 hours. Plates were incubated at room temperature with biotin-conjugated secondary antibody for 1 hour, followed by Avidin-HPR and detection with a TMB Substrate Set (BioLegend). Plates were washed 4 times with PBS containing 0.05% Tween-20 between each step.

### 2.8 Metabolic analysis

The Seahorse XFe96 Analyzer was used to measure mitochondrial respiration and glycolysis simultaneously following a modified protocol. Briefly, BMDMs were plated at a density of 6×10^4^ cells/well in 80μL of culture medium in an Agilent Seahorse XF96 Cell Culture Microplate. After 3 hours, an additional 80 μL of medium was added containing permethrin (10 μM final) and/or pyridostigmine bromide (100 ng/mL final) as indicated. After incubating overnight, an additional 40 μL culture medium containing LPS (200 ng/μL final) was added as indicated. Following a 6 hr incubation, cells were washed and media was replaced with XF assay medium (Base Medium Minimal DMEM lacking glucose, glutamine, and bicarbonate, supplemented with 2 mM Ala-Gln, pH 7.4) prior to analysis. Oxygen consumption rate (OCR) and extracellular acidification rate (ECAR) were measured after sequential addition of glucose 25 mM; oligomycin (1.5 uM); FCCP (1.5 μM) + sodium pyruvate (1mM) and; antimycin A (833 nM) + rotenone (833 nM). Beginning the run with glucose omitted from the medium allows both respirator and glycolytic parameters to be determined in the same run. Respiratory and glycolytic parameters were calculated using the equations outlined in the Seahorse XF Mito Stress Test and Glycolysis Stress Test manuals.

### 2.9 Statistical analyses

All p-values were calculated using two-tailed, unpaired, Student’s t-test, one-way ANOVA with Tukey’s multiple comparison, or the two-stage step-up method to yield q values that have been corrected for the false discovery rate in Excel or GraphPad Prism as indicated in figure legends.

## 3. Results

### 3.1. Female GWIC-exposed mice display more pronounced impairments in cognitive function and mood than male GWIC-exposed mice

To establish GWIC-dependent cognitive symptoms in our model, we employed a battery of tests to characterize neurobehavioral changes following GWIC exposure. A randomly selected subset of GWIC- and DMSO-treated male and female C57BL/6 wild-type (WT) mice were tested for the hippocampal-dependent ability to perceive changes in the environment using an object location test (OLT)^27^. Notably, at both 5 months and 12 months post-exposure, female GWIC-exposed mice showed impairment, failing to show a statistically significant affinity for the object in a novel location during the testing phase (Figure 1A). Male GWIC-exposed mice did not display this impairment, nor did DMSO-exposed mice, regardless of sex. The same cohorts were tested for pattern separation ability using the pattern separation test (PST), which relies on the integrity of the dentate gyrus of the hippocampus^27^. As observed in the OLT, female GWIC-exposed mice showed impairment in the testing phase, while male GWIC-exposed mice and DMSO-exposed mice of both sexes showed a statistically significant increase in affinity for the novel object in pattern 2, reflecting pattern separation ability (Figure 1B). For both tests, control data for total exploration time, total distance moved, and velocity of movement in the testing phase were similar between testing groups (Supplemental Figure 1), indicating that variable object exploration or motor deficits do not explain the differences in test performance. Rather, these results indicate an impairment in hippocampal function in female, but not male, GWIC-exposed mice at 5 and 12 months post-exposure.

We observed similar sex-differences in tests of mood function following GWIC-exposure. At 5 months post-exposure, we used the elevated plus maze test (EPMT) (Supplemental Figure 2A) to determine unconditioned response to a potentially dangerous environment and anxiety-related behavior^28,29^. We observed a trend in which female, not male, GWIC-exposed mice spent less time in open maze arms (Supplemental Figure 2B) and more time in closed maze arms or the center (Supplemental Figure 2D), consistent with anxiety-related behaviors. Calculation of the ratio of visits to the closed arms or center region of the maze versus visits to the open arms revealed a statistically significant increase in preference for the closed arms or center of the maze in GWIC-exposed female mice, but not GWIC-exposed male mice, as compared to DMSO-exposed counterparts (Supplemental Figure 2C). As with the OLT and PST, control data were similar among all cohorts (Supplemental Figure 2E), ruling out motor deficits as the underlying reason for differences in EPMT performance. These results suggest that an increase in anxiety-related behavior is an additional sex-dependent neurobehavioral change following GWIC-exposure.

### 3.2. Female and male GWIC-exposed mice display similarly impaired object recognition memory

Cohorts of mice also were tested for object recognition memory using the novel object recognition test (NORT). Both male and female GWIC-exposed mice failed to show an affinity for the novel object during the testing phase of the NORT, indicating impaired recognition memory. DMSO-exposed mice, regardless of sex, spent a significantly greater amount of object exploration time with the novel object during the testing phase (Figure 1C). As with the OLT and PST tests, control data were comparable among treatment groups (Supplemental Figure 1), indicating that variable object exploration or motor deficits do not explain the differences in test performance.

While both male and female GWIC-exposed mice display impairment on the NORT, which depends primarily upon the integrity of the perirhinal cortex region of the cerebral cortex and partially the hippocampus^27^, only female GWIC-exposed mice display impairment on two tests (OLT, PST) that rely on hippocampal function. This suggests that female GWIC-exposed mice may exhibit increased hippocampal pathology, which may include mitochondrial alterations and neuroinflammation, compared to GWIC-exposed male mice.

### 3.3. GWIC-exposed mice display sex-dependent alterations in expression of mitochondrial proteins in the hippocampus and cerebral cortex

To evaluate mitochondrial changes in the brains of these male and female GWIC-exposed mice, we performed comparative expression analysis of regions required for performance of these neurocognitive tests: the hippocampus, required for the OLT, NORT, and PST, and the cerebral cortex, required for the NORT. Following completion of behavioral tests at 12 months, brains were microdissected to isolate the hippocampus and cerebral cortex (CC), and protein extracts generated were subjected to Western blotting and densitometry quantification.

Examination of an array of mitochondrial proteins revealed widespread rewiring of mitochondrial protein expression, particularly enzymes involved in oxidative phosphorylation complexes I-V (OXPHOS CI-CV). Interestingly, the directionality of the expression changes varied between male and female mice, with the expression of several OXPHOS proteins (NDUFB8, UQCRC2, and MT-COI) significantly increasing in the hippocampi of GWIC-exposed female mice (Figure 2A) but trending towards a decrease in the hippocampi of GWIC-exposed male mice (Figure 2B). Similarly, moderate reductions in abundance of MnSOD (mitochondrial superoxide dismutase that generates hydrogen peroxide), OGDH (TCA cycle enzyme), and TFAM (mtDNA packaging protein) were observed in female, but not male, GWIC-exposed hippocampi. Notably, this observation of more pronounced changes in mitochondrial expression within the hippocampus of female GWIC-exposed mice parallels our earlier finding that GWIC-exposed female mice show greater impairment in hippocampal-dependent neurocognitive tests.

**Figure 2.**
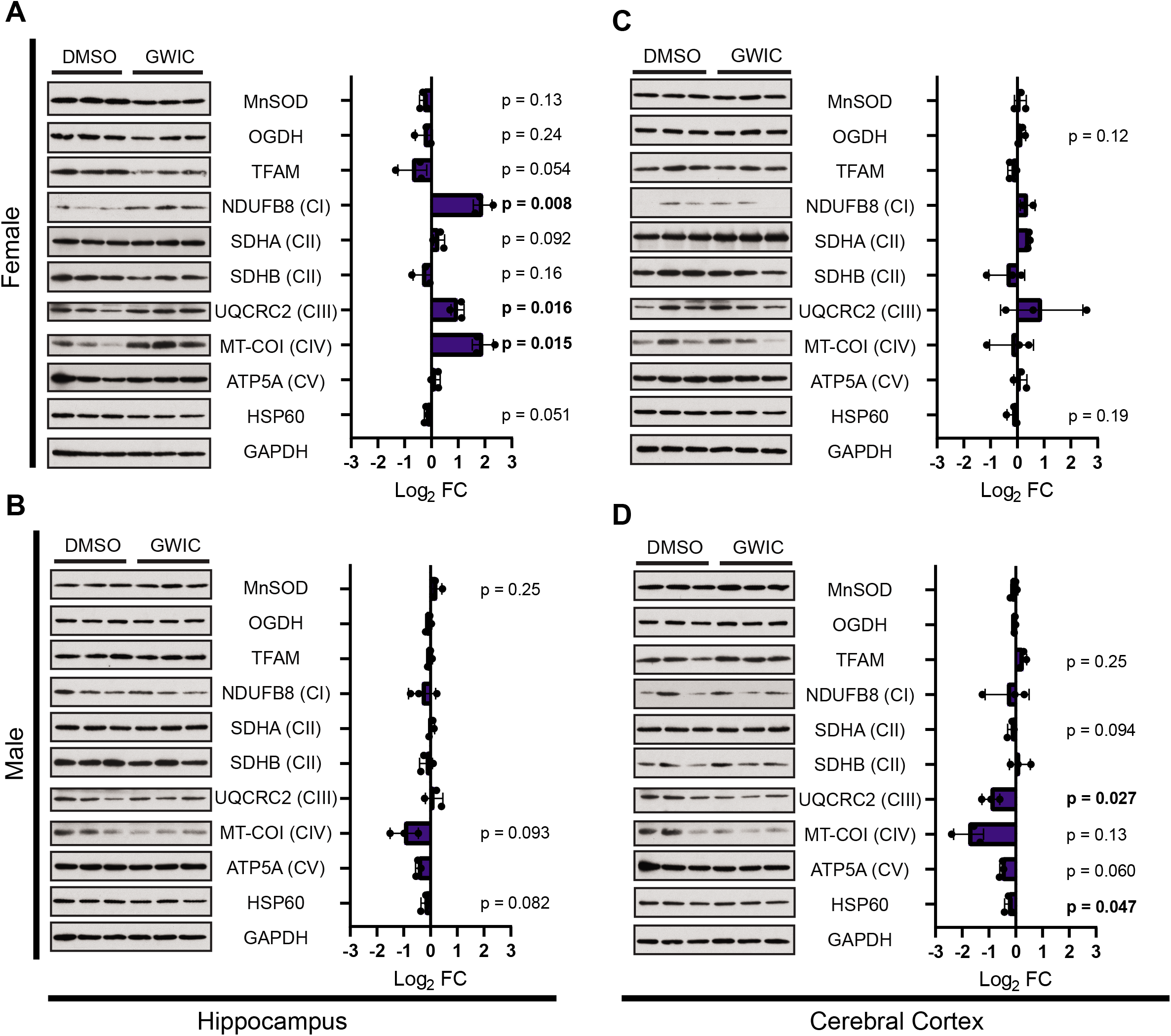
Exposure of C57BL/6 wild-type mice to GWIC results in sex-dependent alterations in expression of mitochondrial proteins in the hippocampus and cerebral cortex. At 12 months post-exposure, tissue extracts were generated from microdissected hippocampi from DMSO- and GWIC-exposed female mice (A) and male mice (B) and from cerebral cortex samples from DMSO- or GWIC-exposed female mice (C) and male mice (D), and immunoblotted for the listed proteins as described in Section 2.5 of materials and methods. Labels in parentheses next to listed proteins designate proteins as part of complex I (CI), complex II (CII), complex III (CIII), complex IV (CIV) or complex V (CV) of the electron transport chain. Densitometry for each protein was performed in ImageJ and normalized to the GAPDH expression level. Bars represent the log_2_ fold change (FC) of biological replicates +/- SD (n=3). Indicated p-values were calculated using a two-tailed t test, and values below 0.05 are shown in bold.

Analysis of extracts from the CC also reveals sex-dependent directionality of expression changes. Although expression of UQCRC2 was largely unchanged in the hippocampi of GWIC-exposed male mice, it is significantly decreased in CC extracts from these animals (Figure 2D). In contrast, UQCRC2 levels in CC extracts isolated from female GWIC-exposed mice show a modest increase (Figure 2C), echoing the varied directionality observed in the hippocampus. Similarly, we observed a trending decrease in expression of MT-COI, ATP5A, and SDHA in CC extracts of male, but not female, mice. Overall, these changes in mitochondrial protein abundance reveal sex-dependent OXPHOS remodeling, which persists for 12 months after exposure to GWIC and tracks with neurocognitive impairments.

### 3.4. Female GWIC-exposed mice exhibit increased neuroinflammation as compared to male GWIC-exposed mice

To assess neuroinflammation after GWIC exposure, perfused brains from female and male mice were fixed, sectioned, and processed for immunohistochemical quantification of glial fibrillary acidic protein (GFAP)-positive astrocytes and ionized calcium-binding adaptor molecular 1 (IBA-1)-positive microglia. Quantification of astrocytes in the dentate gyrus (DG), CA1, and CA3 regions of the hippocampus (Figure 3) revealed that female GWIC-exposed mice displayed significantly increased GFAP+ structures in all three hippocampal regions as compared to female DMSO-exposed mice. Male GWIC-exposed mice show significantly increased GFAP+ structures in the DG region and CA3 region only. This observation, which indicates more widespread hippocampal inflammation in female GWIC-exposed mice as compared to male GWIC-exposed mice, is consistent with our findings that female GWIC-exposed mice display increased impairment on hippocampal-dependent neurobehavioral tests.

**Figure 3.**
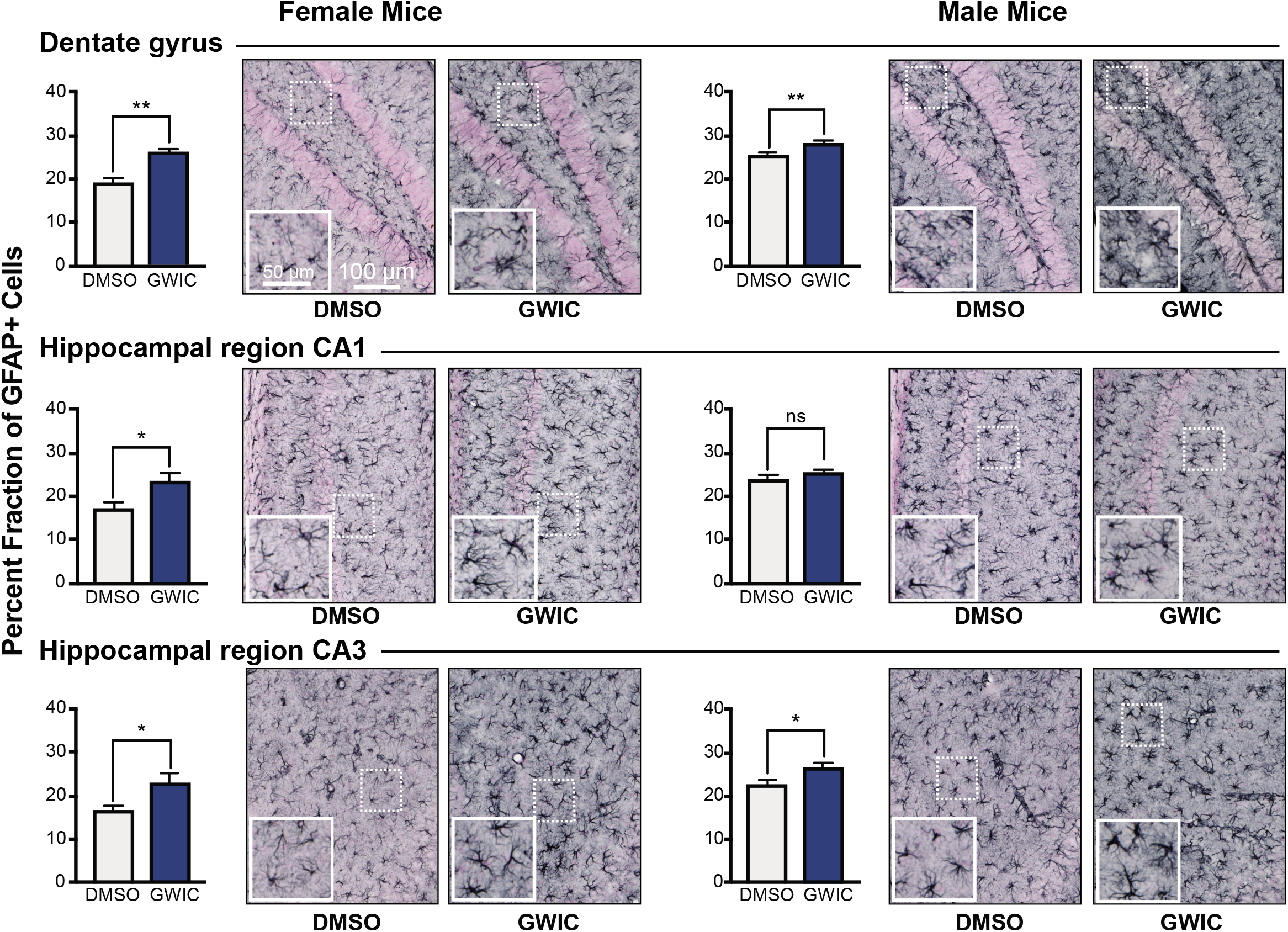
Increased astrocyte hypertrophy is observed in three hippocampal regions of female GWIC-exposed mice and in two hippocampal regions of male GWIC-exposed mice. We measured percent fraction of glial-fibrillary acidic protein-expressing (GFAP+) cells within three regions of the hippocampus: the dentate gyrus (DG), CA1, and CA3. Images show representative distribution and morphology of GFAP+ astrocytes in these three regions in male and female GWIC-exposed and DMSO-exposed mice (scale bar = 100 μM). Insets show magnified views (scale bar = 50 μM); adjacent bar graphs show quantification of percent fraction GFAP+ for each cohort). Error bars represent the mean of biological replicates +/- SEM (n=6). Indicated p-values were calculated using two-tailed, unpaired, Student’s t-test.. *p < 0.05; **p < 0.01; ns, not significant.

A study by Zakirova and colleagues of neuroinflammation in this murine model of GWIC-exposure have likewise observed significant increases of astrocyte hypertrophy in the dentate gyrus of the hippocampus, as measured by GFAP quantification. Notably, this study, which exclusively used male mice, did not observe a significant increase in activated IBA-1+ microglia following GWIC-exposure^24^. In contrast, when we performed blinded morphological scoring of IBA-1+ cells with the DG, CA1, and CA3 of hippocampi from GWIC- and DMSO-exposed male and female mice, we observed a significant increase in activated microglia within the DG of female, but not male, GWIC-exposed mice as compared to DMSO-exposed mice (Supplemental Figure 7A). Significant increases were not observed in the CA1 or CA3 hippocampal regions of female or male GWIC-exposed mice, and male GWIC-exposed mice displayed a significant reduction in IBA-1+ cells with an activated morphology within the CA1 region, consistent with decreases in percent fraction of IBA-1+ cells observed by Zakirova *et al*. Our findings therefore suggest a novel sex-dependent increase in activated microglia within the DG region of the hippocampus in this model of GWIC-exposure.

### 3.5. GWIC-exposure of cells results in metabolic alterations, enhanced expression of interferon-stimulated genes, and increased production of inflammatory cytokines

Our findings are consistent with the notion that GWIC exposure causes lasting mitochondrial rewiring and neuroinflammation. However, a direct effect of GWIC on metabolism and/or innate immune signaling has not been documented. Therefore, to characterize the immediate impact of GWIC on mitochondrial function and cell-intrinsic innate immune responses, we performed a series of acute, in vitro experiments using bone marrow-derived macrophages (BMDMs). While not derived from the brain, BMDMs provide a robust cellular platform to evaluate mitochondrial function, innate immune responses, and the dynamic interplay between these pathways^30^. To first evaluate GWIC-dependent changes in oxygen consumption and glycolysis, BMDMs were treated overnight with permethrin (Per) alone, pyridostigmine bromide (PB) alone, or, as in *in vivo* experiments, both of these GWIC (Per-PB), then subjected to analysis of mitochondrial respiration using a Seahorse XF96 Analyzer. Per or Per-PB exposed BMDMs exhibited significantly increased basal mitochondrial oxygen consumption (OCR), with unchanged maximal respiration (Figure 4A), while treatment with PB only failed to increase basal respiration and resulted in significantly diminished maximal respiration. Treatment with Per, PB, or Per-PB decreased spare respiratory capacity, accompanied by an increase in proton leak for Per- and Per-PB-treated BMDMs. Overall, these data indicate that acute exposure to GWIC induces changes to mitochondrial OXPHOS and are consistent with the in vivo findings described in Section 3.3. Additional changes were observed when BMDMs were treated with bacterial lipopolysaccharide (LPS, an agonist of the innate immune sensor Toll-like receptor 4 (TLR4)) in combination with GWIC. Per or Per-PB pretreatment followed by LPS resulted in an increase in basal respiration, as seen in mock-treated cells, as well as a significant increase in maximal respiration (Figure 4B). Interestingly, LPS treatment resulted in significantly increased spare respiratory capacity upon exposure to all three GWIC combinations, though this could be due to the increase in maximal respiration. Following LPS treatment, proton leak was increased only in Per cells. Additional studies analyzing a surrogate marker of glycolysis, the extracellular acidification rate (ECAR), revealed that while glycolysis was largely unchanged following GWIC-exposure alone (Supplemental Figure 3B), treatment with GWIC plus LPS resulted in cells becoming more glycolytic (Supplemental Figure 3C), consistent with the immunometabolic changes typically seen in inflammatory M1 macrophages.

**Figure 4.**
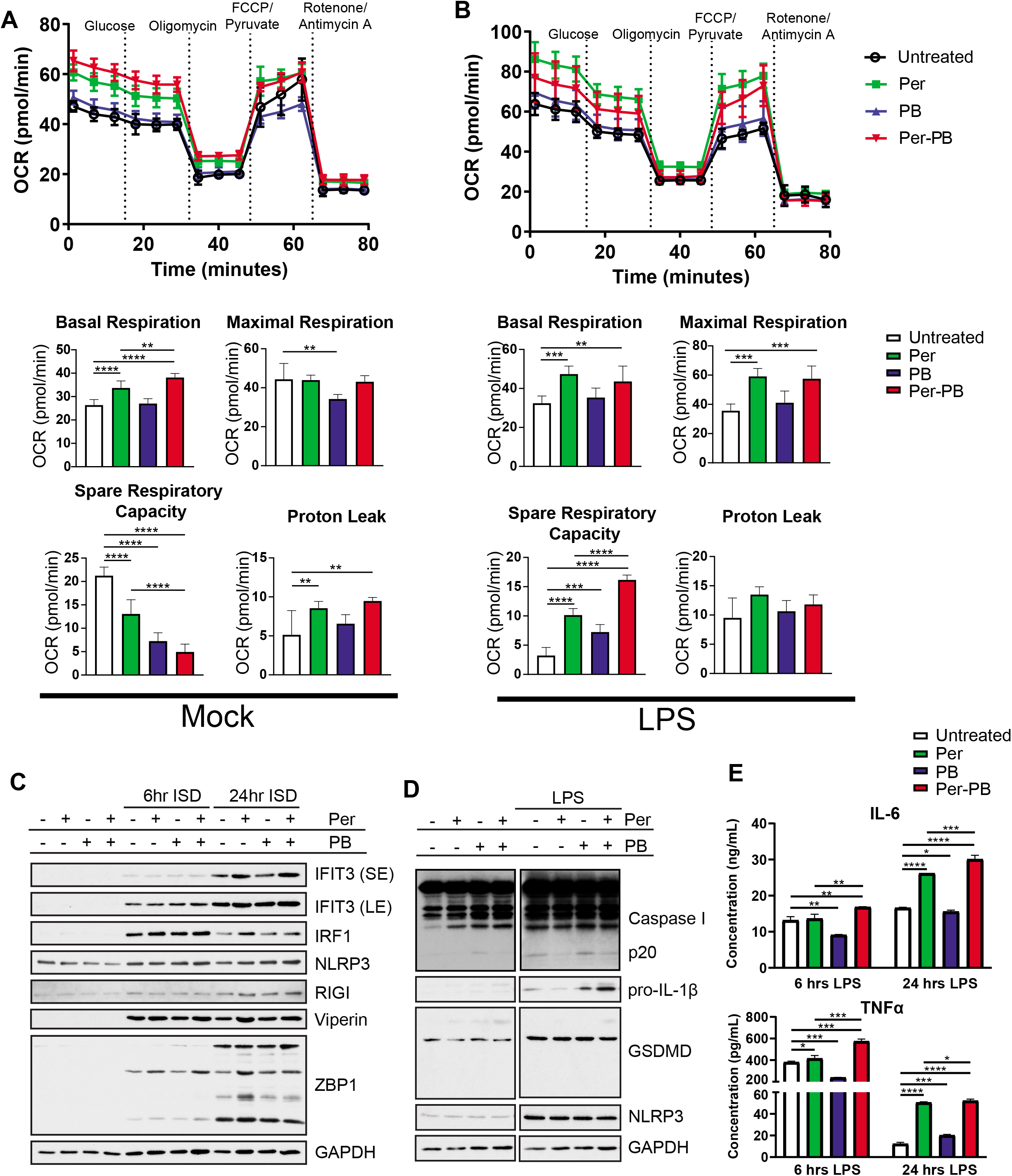
GWIC exposure alters mitochondrial respiration and potentiate inflammatory responses in BMDMs. (A) and (B) Oxygen consumption rate (OCR) in BMDMs treated with GWIC. BMDMs were treated overnight with GWIC as indicated. Cells were treated with (A) Seahorse medium (Mock) or (B) 200 ng/mL LPS (LPS) for 6 hours, then respiration was measured on a Seahorse XF96 Analyzer using a modified protocol as described in Section 2.8 of materials and methods. (C) ISG expression in BMDMs treated with GWIC and transfected with ISD. BMDMs were treated overnight with GWIC as indicated, then transfected with 1 μg ISD for 6 or 24 hours. Cells extracts were blotted for indicated proteins as in Figure 2. (D) Inflammasome activation in BMDMs treated with GWIC. BMDMs were treated overnight with GWIC as indicated, then treated with 20 ng/mL LPS for 6 or 24 hours. Cells extracts were blotted for indicated proteins as in Figure 2. (E) ELISA of inflammatory cytokines in BMDMs treated as in (D). Data shown is representative of three separate experiments. Error bars represent the mean of technical replicates +/- SD (n=5 for (A) and (B) and n=2 for (E)). Significance values for pairwise comparisons were calculated using the two-stage step-up method to yield q values that have been corrected for the false discovery rate. * q < 0.05; ** q < 0.01; *** q < 0.001; **** q < 0.0001.

Prior work has shown that systemic LPS-induced TLR4 activation contributes to GWIC-related neuroinflammation^31^. To examine the direct effects of GWIC exposure on innate immune signaling, we first measured expression of induced proteins following LPS-stimulation of BMDMs that had been pre-treated with Per, PB, or Per-PB. We observed modest increases in expression of the interferon regulatory factor IRF1 in BMDMs exposed to LPS plus Per and/or Per-PB, as well as modestly elevated levels of the interferon stimulated gene (ISG) IFIT3 after 6 hours of LPS plus PB and Per-PB exposure (Supplemental Figure 3A). Additional work in BMDMs revealed synergy between the STING innate immune pathway, as transfection of interferon stimulatory DNA (ISD) to activate STING after pretreatment with GWIC resulted in modestly elevated ISG expression (Figure 4C). Moreover, we noted an increase in pro-IL-1β cytokine abundance after LPS plus GWIC (Figure 4D). In fact, PB increased Caspase I cleavage to the active p20 isoform in both untreated and LPS-treated cells, a process that requires activation of cytosolic inflammasomes such as NLRP3. Finally, ELISA analysis of BMDMs revealed that GWIC exposure synergized with LPS stimulation and resulted in increased proinflammatory cytokine production: Per or Per-PB treatment led to increased production of both IL-6 and TNF-α after 24 hours of LPS treatment (Figure 4E). Interestingly, though PB alone inhibited IL-6 and TNF-α production, it acted synergistically with Per and enhanced IL-6 and TNF-α production over Per treatment alone. Overall, our results reveal that GWIC induce immunometabolic rewiring of OXPHOS and glycolysis in BMDMs and potentiate inflammatory responses to innate immune stimulation.

### 3.6. Ablation of STING or NLRP3 signaling improves neurocognitive deficits and neuroinflammation and blunts changes in mitochondrial protein expression following GWIC-exposure

Our in vivo experiments using a mouse model of GWI have demonstrated neurocognitive deficits, changes in expression of mitochondrial proteins, and hippocampal inflammation caused by GWIC, with more severe symptoms observed in female mice. Our *in vitro* experiments using BMDMs revealed metabolic changes and heightened innate immune responses following GWIC treatment, including increased production of ISGs and the proinflammatory cytokines IL-1β, IL-6, and TNFα. Notably, BMDM experiments suggested potentiated TLR4, STING, and NLRP3 innate immune signaling in response to acute GWIC exposure.

Stress, infection, or injury can cause MD, which leads to the release of mitochondrial constituents such as mtDNA, mitochondrial lipids (cardiolipin), *N*-formyl peptides, mitochondrial metabolites (succinate), and high concentrations of ATP^21^. These so-called damage-associated molecular patterns (DAMPs) then trigger the innate immune system via several paths. mtDNA can be detected by the enzyme cGAS, which then generates the cyclic dinucleotide cGAMP, a second messenger that binds and activates STING^32^ for expression of type I interferons and ISGs^33^. Likewise, over-production of ROS, mtDAMP release, and altered mitochondrial dynamics due to MD engage the NLRP3 inflammasome, triggering processing of pro-interleukin-1 beta (pro-IL-1β) into IL-1 β^34,35^. Both STING and NLRP3 are therefore key mechanisms by which MD and mtDAMPs can engage the innate immune system, and the increases in ISGs and IL-1β observed in Figure 4C, Figure 4D, and Supplemental Figure 3A implicate both pathways downstream of GWIC.

To interrogate the contribution of STING and NLRP3 signaling to neurobehavioral changes, neuroinflammation, and mitochondrial rewiring following GWIC exposure, cohorts of STING^gt/gt^ and NLRP3^−/−^ mice on a C57BL/6 background were exposed to DMSO or GWIC in parallel to wild-type C57BL/6 (WT) mice. STING^gt/gt^ mice are homozygous for an I199N missense mutant allele in the *Sting* gene and express no STING protein^36^, while NLRP3^−/−^ mice are homozygous for a neo cassette in the coding region of *Nlrp3* and are therefore deficient in the NLRP3 inflammasome^37^.

Neurobehavioral tests of STING^gt/gt^ mice revealed that observed impairments of GWIC-exposed mice are STING-dependent. In contrast to WT GWIC-exposed cohorts, GWIC-treated STING^gt/gt^ female mice demonstrated rescue of object location memory by successfully completing the OLT at both 5 months post-exposure (Figure 5A) and at 12 months post-exposure (Supplemental Figure 4A). Similarly, GWIC-treated STING^gt/gt^ female mice at 5 months post-exposure (Figure 5A) and GWIC-treated STING^gt/gt^ male mice at 12 months post-exposure (Supplemental Figure 4B) were unimpaired in the NORT. These findings suggest that STING signaling may contribute to impairment of both hippocampal function in female mice and cerebral cortex function in male and female mice following GWIC exposure.

**Figure 5.**
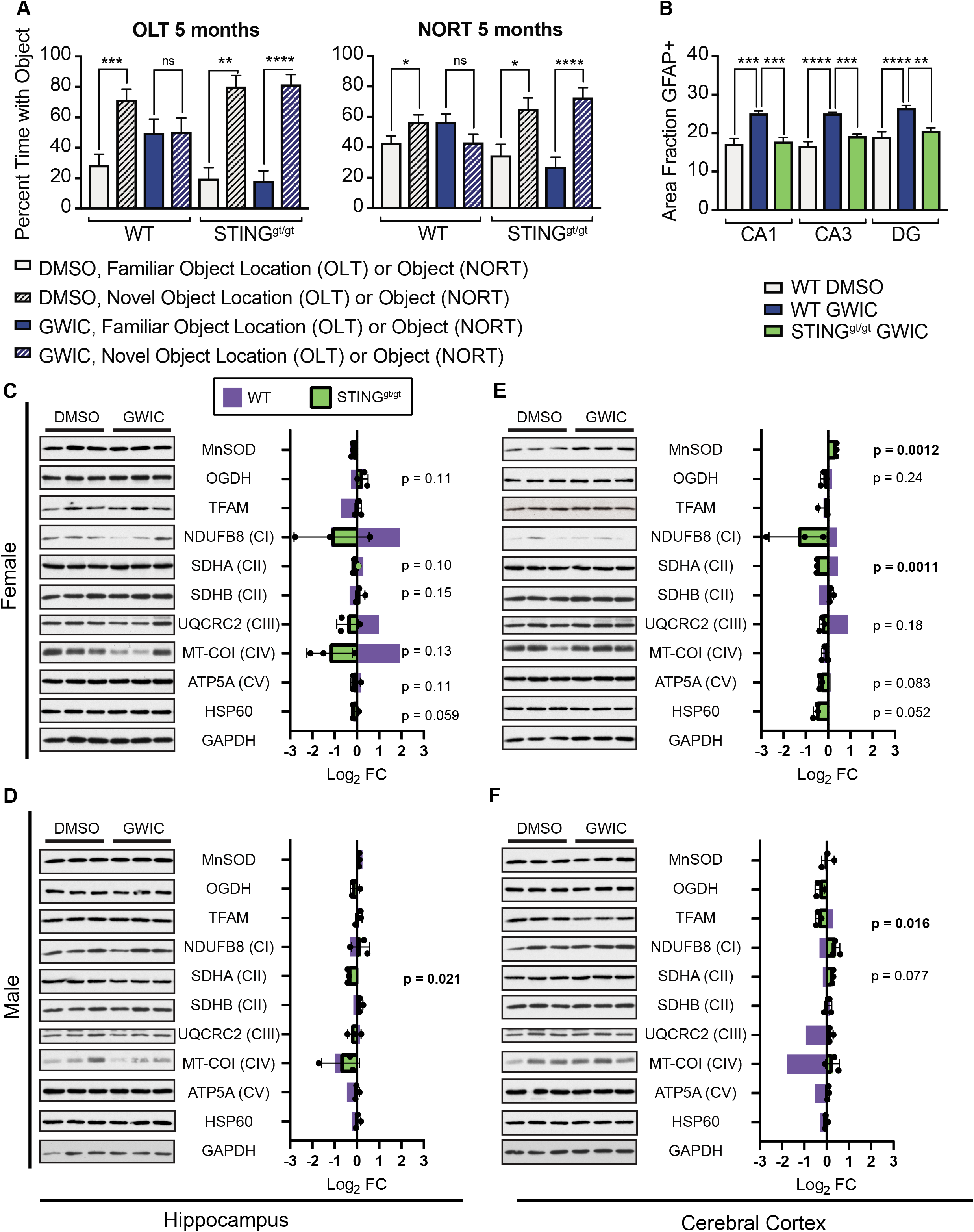
Loss of STING attenuates GWIC-related neurobehavioral changes, neuroinflammation, and mitochondrial rewiring. (A) A comparison of OLT and NORT results in DMSO- or GWIC-exposed WT and STING^gt/gt^ C57BL/6 female mice. Error bars represent the mean of biological replicates +/- SEM (OLT: n=3 for DMSO control mice, n=6 for GWIC-exposed mice; NORT; n=4 for DMSO control mice, n=8 for GWIC-exposed mice). (B) Comparison of percent fraction of GFAP+ cells within the dentate gyrus (DG), CA1, and CA3 of DMSO- or GWIC-exposed WT female mice and GWIC-exposed STING^gt/gt^ female mice. Indicated p-values for (A) and (B) were calculated using two-tailed, unpaired, Student’s t-test. *p < 0.05; **p < 0.01; ***p < 0.001; ****p < 0.0001; ns, not significant. (C) Hippocampal extracts from DMSO- and GWIC-exposed female (C) and male (D) WT and STING^gt/gt^ mice and cerebral cortex extracts from DMSO- and GWIC-exposed female (E) and male (F) WT and STING^gt/gt^ mice 12 months post-exposure were generated and blotted as described in Figure 2. Green bars represent the log_2_ fold change (FC) of biological replicates +/- SD (n=3) of STING^gt/gt^ extracts, with each replicate represented by a black dot. Blue bars in overlay represent the log_2_ fold change (FC) of biological replicates (n=3) of WT samples from Figure 2, for sake of comparison. Densitometry for each protein was performed in ImageJ and normalized to the GAPDH expression level. Indicated p-values, which refer to STING^gt/gt^ samples, were calculated using a two-tailed t test, and values below 0.05 are shown in bold.

Interestingly, neurobehavioral tests of NLRP3^−/−^ mice suggest that neurobehavioral changes due to GWIC exposure are only partially NLRP3-dependent. Like GWIC-treated female WT mice, female NLRP3^−/−^ mice display impaired object location recognition at 5 months (Figure 6A) and 12 months (Supplemental Figure 4A) post-exposure, indicating that NLRP3 signaling is not important for the development of this neurocognitive change. Additionally, although female NLRP3^−/−^ mice at 5 months post-exposure show restored novel object recognition (Figure 6A), this restoration was lost at 12 months post-exposure (Supplemental Figure 4B).

**Figure 6.**
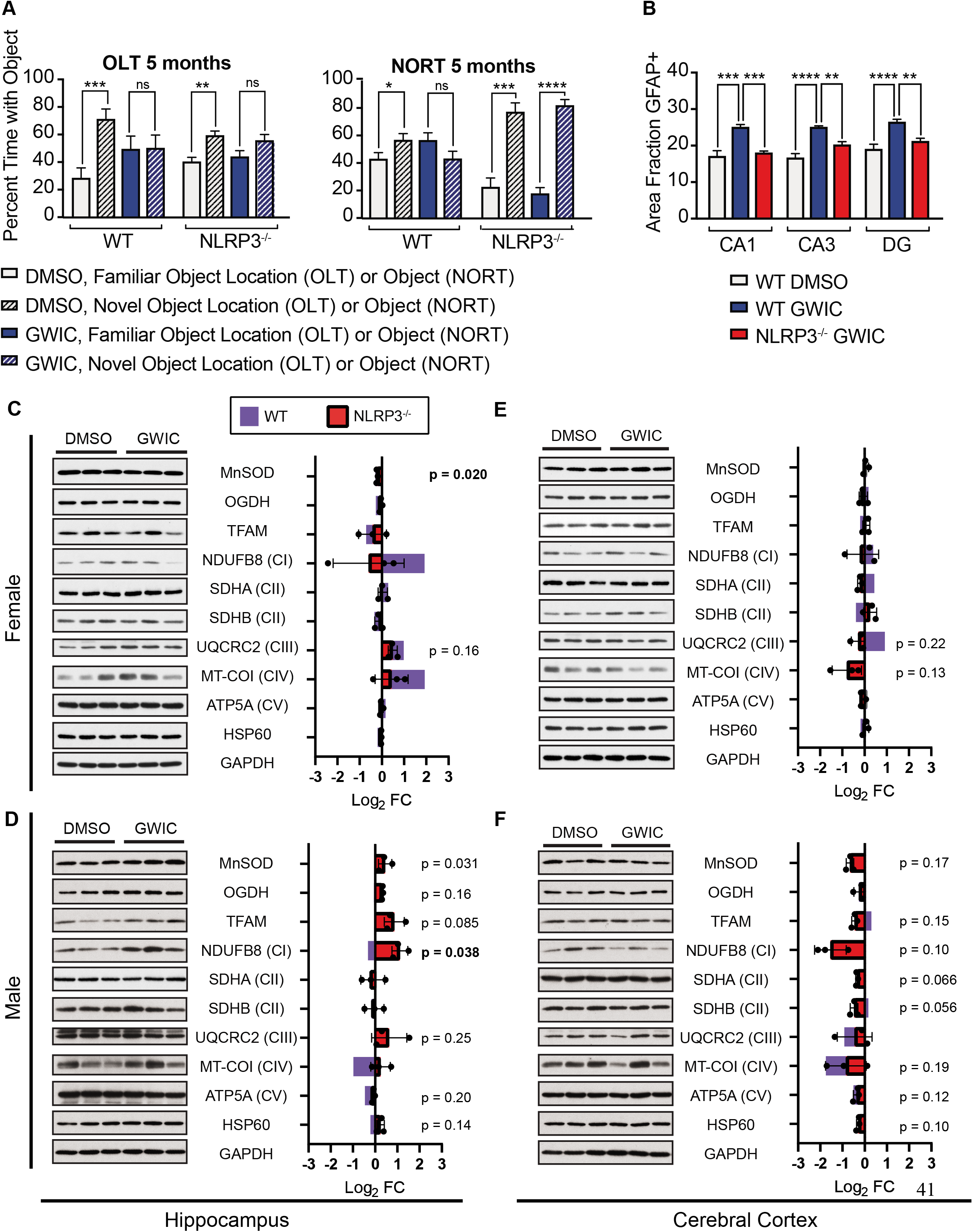
Loss of NLRP3 (partly) attenuates GWIC-related neurobehavioral changes, neuroinflammation, and mitochondrial rewiring. **(A)** A comparison of OLT and NORT results in DMSO- or GWIC-exposed WT and NLRP3^−/−^ C57BL/6 female mice. Error bars represent the mean of biological replicates +/- SEM (OLT: n=5 for DMSO control mice, n=10 for GWIC-exposed mice; NORT; n=5 for DMSO control mice, n=9 for GWIC-exposed mice). (B) Comparison of percent fraction of GFAP+ cells within the dentate gyrus (DG), CA1, and CA3 of DMSO- or GWIC-exposed WT female mice and GWIC-exposed NLRP3^−/−^ female mice. Indicated p-values for (A) and (B) were calculated using two-tailed, unpaired, Student’s t-test. *p < 0.05; **p < 0.01; ***p < 0.001; ****p < 0.0001. (C) Hippocampal extracts from DMSO- and GWIC-exposed female (C) and male (D) WT and NLRP3^−/−^ mice and cerebral cortex extracts from DMSO- and GWIC-exposed female (E) and male (F) WT and NLRP3^−/−^ mice 12 months post-exposure were generated and blotted as described in Figure 2. Red bars represent the log_2_ fold change (FC) of biological replicates +/- SD (n=3) of NLRP3^−/−^ extracts, with each replicate represented by black dots. Blue bars in overlay represent the log_2_ fold change (FC) of biological replicates (n=3) of WT samples from Figure 2, for sake of comparison. Densitometry for each protein was performed in ImageJ and normalized to the GAPDH expression level. Indicated p-values, which refer to NLRP3^−/−^ samples, were calculated using a two-tailed t test, and values below 0.05 are shown in bold.

Ablation of STING signaling and the NLRP3 inflammasome appears to improve mood function in female mice following GWIC-exposure. While WT GWIC-exposed female mice displayed a statistically significant preference for the closed arms or center of the maze during the EPMT, STING^gt/gt^ and NLRP3^−/−^ female mice fail to show this preference (Supplemental Figure 4D). STING^gt/gt^ and NLRP3^−/−^ GWIC-exposed male mice behave similarly to WT counterparts, without an observed preference (Supplemental Figure 4D).

The results of the experiments described above demonstrate that STING and NLRP3 contribute to GWI-dependent neurocognitive changes to a varying degree. Additional neurobehavioral tests were more difficult to interpret at the inter-trial intervals employed in the study. For example, both DMSO- and GWIC-treated STING^gt/gt^ female mice are deficient in novel object recognition at 12 months post-exposure, indicating non-treatment specific, aging-related neurobehavioral changes in these mice (Supplemental Figure 4B). Similarly, STING^gt/gt^ and NLRP3^−/−^ female and male mice showed impaired pattern separation ability, regardless of treatment, at 5 months post-exposure (Supplemental Figure 4C). Because of this, we cannot draw conclusions from 12 month post-exposure NORT tests in female mice or 5 month post-exposure PST tests in female or male mice. While STING signaling or the NLRP3 inflammasome may in fact impact neurobehavioral changes as measured by these assays, changes to the DMSO-treatment control mice make conclusions impossible.

Other tests were also inconclusive. The validity of NORT testing of GWIC-treated STING^gt/gt^ male mice at 5 months and NLRP3^−/−^ male mice at 5 and 12 months is uncertain due to the failure of DMSO-treated control cohorts to spend significantly increased time with the novel object (Supplemental Figure 4B). In the case of the DMSO-treated STING^gt/gt^ mice at 5 months post-exposure, controls measurements indicated decreased movement and object exploration (Supplemental Figure 5). However, for all other groups and tests, control data indicate that variable object exploration or motor deficits do not explain test performance (Supplemental Figure 5).

In the WT GWIC-exposed female mice, deficiencies in neurobehavioral tests correlated with increased astrocyte hypertrophy in three regions of the hippocampus. In contrast, both STING^gt/gt^ and NLRP3^−/−^ female mice failed to show increased astrocyte hypertrophy in the DG (Figure 5B, Figure 6B) or the CA1 and CA3 regions (Supplemental Figure 6A, 6B) as compared to DMSO-treated WT controls. Expression of GFAP in the hippocampi of DMSO-treated WT, STING^gt/gt^, and NLRP3^−/−^ were comparable (Supplemental Figure 6C), suggesting that this observation is due to lessened GWIC-induced neuroinflammation in the absence of STING or NLRP3. In addition, analysis of IBA-1+ cells within the hippocampus also revealed a decrease in activated microglia within the DG of STING^gt/gt^, and NLRP3^−/−^ GWIC-exposed female mice (Supplemental Figure 7A, 7B) as compared to WT GWIC-exposed female mice.

STING and NLRP3 therefore play a role in driving neurocognitive changes and neuroinflammation following GWIC exposure. To assess the impact on mitochondrial expression changes, extracts of the hippocampi (Figure 5C and 5E, Figure 6C and 6E) and cerebral cortices (Figure 5D and 5F, Figure 6D and 6F) of DMSO- and GWIC-exposed STING^gt/gt^ and NLRP3^−/−^ mice were analyzed as in Figure 2. Hippocampal extracts from both STING^gt/gt^ (Figure 5C) and NLRP3^−/−^ female mice (Figure 6C) no longer show the significant increase in NDUFB8 observed in WT female mice, and STING^gt/gt^ female mice also show a downward trend in UQCRC2 expression and a significant decrease in MT-COI, in contrast to the significant upregulation of both in WT female mice. We also observed an upward trend of expression for UQCRC2, MT-COI, ATP5A, and SDHA in the CC of STING^gt/gt^ males (Figure 5F), which were all downregulated in WT males. Overall, expression analysis of STING^gt/gt^ and NLRP3^−/−^ brains echoes other findings with these mice: the absence of these signaling hubs reduces cognitive impairment, neuroinflammation, and mitochondrial expression changes, with findings indicating a more significant role for STING than for NLRP3.

## 4. Discussion

Using an established murine model of GWI^22,29,24^, we demonstrated that GWIC-exposed female mice display impaired object location memory and novel object recognition at 5 and 12 months post-exposure, and impaired pattern separation at 5 months post-exposure. These results show that female mice have more severe GWIC-dependent neurocognitive impairment than males, which in our studies showed impairment only in novel object recognition and no anxiety-related behavior as measured by the EPMT. A previous study examining both female and male C57BL/6 mice also demonstrated increased anxiety in female mice, as measured by the open field test (OFT)^38^. However, the majority of GWI rodent model studies have used male mice only, using tests including the Barnes maze^24,29,7,39^, forced swim test^39,40^, and OFT^38,35,26^ to reveal cognitive impairment, fatigue, and anxiety-related behavior. We tested both female and male GWIC-exposed mice using the OLT, NORT, and PST, which has been used in GWI rat models to show impairment in male animals with demonstrated hippocampal inflammation^27,42,43,44^. We also used the EPMT, which has shown disinhibition type behaviors in GWIC-exposed male mice^29^ that we did not observe, likely due to differences in time post-exposure. This study is therefore the first time female GWIC-exposed mice have been evaluated for neurocognitive impairment using this battery of tests. While women represented a relatively small percentage (7%) of the active duty population during PGW-1^45^, chronic GWI conditions differ in female and male PGW-1 veterans^46^, with some studies showing female veterans PGW-1 as more likely to meet the clinical criteria of GWI and to display heightened, longer-lasting GWI symptoms than male veterans^47,48^. Therefore, there is a critical need to better understand sex differences in GWI pathology. The more significant neurocognitive impairment that we observed in female mice makes a strong argument for inclusion of both sexes in future pre-clinical GWI studies involving rodents.

In addition to cognitive impairment, we observed astrocyte hypertrophy within all three examined regions (dentate gyrus, CA1, and CA3) of the hippocampi of GWIC-exposed female mice, as compared to only two regions (DG and CA3) in male mice. Comparison of DMSO-exposed female and male mice revealed that control males appear to have heightened astrocyte hypertrophy as compared to control females, which may explain the diminished GWIC-dependent increase. We also observed an increase in IBA-1+ microglia with an activated morphology in the dentate gyri of GWIC-exposed female mice, with a decrease observed in GWIC-exposed male mice, consistent with a previous study^29^. Overall, this novel observation of more severe hippocampal inflammation observed in GWIC-exposed female mice further emphasizes the value of using both male and female mice for future GWI study.

Our analysis of expression of mitochondrial proteins in the hippocampi of GWIC-exposed mice adds to the growing body of evidence that MD is an important factor in the pathology of GWI. MD^49^ and mtDNA lesions^50^ have been documented in GWI patients, and animal studies analyzing expression of nuclear and mitochondrially encoded OXPHOS proteins from whole brain extracts have demonstrated widespread expression changes in mice^51^ and rats^3^. We observed decreases in OXPHOS proteins in male mice exposed to GWICs in both the hippocampus and peripheral cortex. In contrast, OXPHOS protein expression in female mice increased in response to GWICs, particularly in the hippocampus, while other mitochondrial proteins including SDHB, MnSOD, TFAM, and OGDH show a modest decrease in expression. This suggests a general rewiring of mitochondrial protein expression, rather than a wholesale decrease in all mitochondrial proteins, that is most severe in hippocampi of female mice.

*In vitro* studies centered around acute treatment of BMDMs provided further evidence of disruptions to mitochondrial metabolism. Of the two GWIC, permethrin was the main driver for alterations to mitochondrial respiration, leading to an increase in basal respiration, loss of spare respiratory capacity, and increase in proton leak. Though PB alone had little effect, it appeared to act synergistically with Per, causing further alterations to basal respiration and spare respiratory capacity in control cells and an increase in spare respiratory capacity in LPS-treated cells. We observed a similar synergistic interaction when we analyzed production of IL-6 and TNFα after LPS stimulation. Per alone induced an increase in cytokine release, while PB alone had little effect; however, treatment with Per and PB together induced an even stronger response. A likely explanation for this observation is that Per and PB are acting through separate but complimentary pathways. BMDMs treated with Per showed increased expression of ISGs than either untreated or PB treated cells after induction with LPS or ISD. Alternatively, BMDMs treated with PB displayed increased expression of pro-IL-1β and increased cleavage of Caspase I. Thought the effect was subtle, this suggests low levels of activation of the NLRP3 inflammasome in response to PB. Consistent with this notion, elevated serum levels of IL-1β and IL-6 have been observed in patients with GWI, and elevated IL-1β, IL-6, and TNFα have been observed in the brains of mice exposed to GWICs^3,43,52–54^.

As in other studies, our analysis of C57BL/6 WT GWIC-exposed mice and of GWIC-treated BMDMs demonstrated contemporaneous MD and inflammation. GWIC-exposure experiments with STING^gt/gt^ and NLRP3^−/−^ mice suggest a link between the two. Loss of STING in female mice restores hippocampal-dependent object location memory at both 5 and 12 months post GWIC exposure and novel object recognition ability at 5 months post-exposure; loss of STING also restores novel object recognition in male GWIC-exposed mice. These data strongly suggest a role for STING in driving neurocognitive changes following GWIC-exposure. Loss of NLRP3 has a subtler effect, restoring novel object recognition in female GWIC-exposed mice at 5 months, but not 12 months, post-exposure. This may indicate a role for NLRP3 in relatively short-term impairment in novel object recognition following GWIC-exposure. These findings are limited by the behavioral tests we employed and by the validity of these tests. As described in Section 3.6, several of the tests of STING^gt/gt^ and NLRP3^−/−^ mice were inconclusive due to impairment observed in control animals or by the failure of cohorts to spend sufficient time in object exploration. However, we were able to draw conclusions from several OLT, NORT, and EPMT tests, which on their own suggest a significant role for STING and a potentially lesser, time-dependent role for NLRP3 in driving neurocognitive impairment.

Analysis of neuroinflammation and expression of mitochondrial proteins within the hippocampi and cerebral cortices of GWIC-exposed mice also support a role for STING and NLRP3 in GWI. Loss of STING or NLRP3 in female mice results in a significant diminishment of astrocyte hypertrophy in all three regions of the hippocampus as compared to GWIC-exposed WT females, as well as a significant decrease in microgliosis within the DG. The rewiring of mitochondrial expression due to GWIC-exposure is also blunted or reversed in STING^gt/gt^ and NLRP3^−/−^ mice. STING and NLRP3 therefore join TLR4^31^ as innate immune sensors linked to GWI-related pathology in murine models. Remaining questions include in which cell types and tissues STING and/or NLRP3 signaling occurs to influence GWI symptoms, as well as the timing of signaling following GWIC exposure. Future study using conditional transgenic models will need to be conducted to address these questions. Comprehensive assessment of these innate immune pathways in patient samples is also needed to confirm whether our findings extend to veterans with GWI. If indeed these pathways are active, targeted intervention using STING and NLRP3 inhibitors may be effective at alleviating some GWI symptoms and improving the healthspan of this population.

Finally, therapeutic targeting of mitochondria to directly improve metabolic function is a viable treatment strategy for GWI. Indeed, veterans with GWI have shown improvements in symptoms in response to Coenzyme Q10 ^55^, an antioxidant and OXPHOS electron carrier known to have beneficial effects on mitochondria, or KPAX002, a combination treatment designed to boost mitochondrial function^56^. Emerging links between mitochondria and the innate immune system raise the possibility that therapeutic approaches to improve mitochondrial health may also serve to decrease inflammation, likely by reducing mtDAMP release/accumulation. Our work implicates two of the main mtDAMP sensing pathways, STING and NLRP3, to GWIC-related inflammation, mitochondrial/metabolic rewiring, and cognitive deficits. Therefore, an intriguing future direction is to evaluate how different mitochondrial therapeutics affect neurocognitive impairment, neuroinflammation, and mitochondrial protein expression within the brain of GWIC-exposed mice to understand the efficacy, mechanisms, and perhaps sex-specific effects, of these treatments.

## Acknowledgements

We thank Sahithi Attaluri for assistance with immunohistochemical analysis and Dr. Alexandra Trott, Associate Director of the TAMU Rodent Preclinical Phenotyping Core, for assistance with behavioral tests. This work was supported, in part, by a grant from NIEHS P30 ESES029067 and by awards W81XWH-17-1-0446 (to A.P.W.), W81XWH-16-1-0480 (to A.K.S.), and W81XWH-17-1-0447 (to A.K.S.) from the Office of the Assistant Secretary of Defense for Health Affairs through the Peer Reviewed Medical Research Programs, Gulf War Illness Research Program. Opinions, interpretations, conclusions, and recommendations are those of the authors and are not necessarily endorsed by the Department of Defense or the United States Government.

---

**Supplemental Figure 1.**
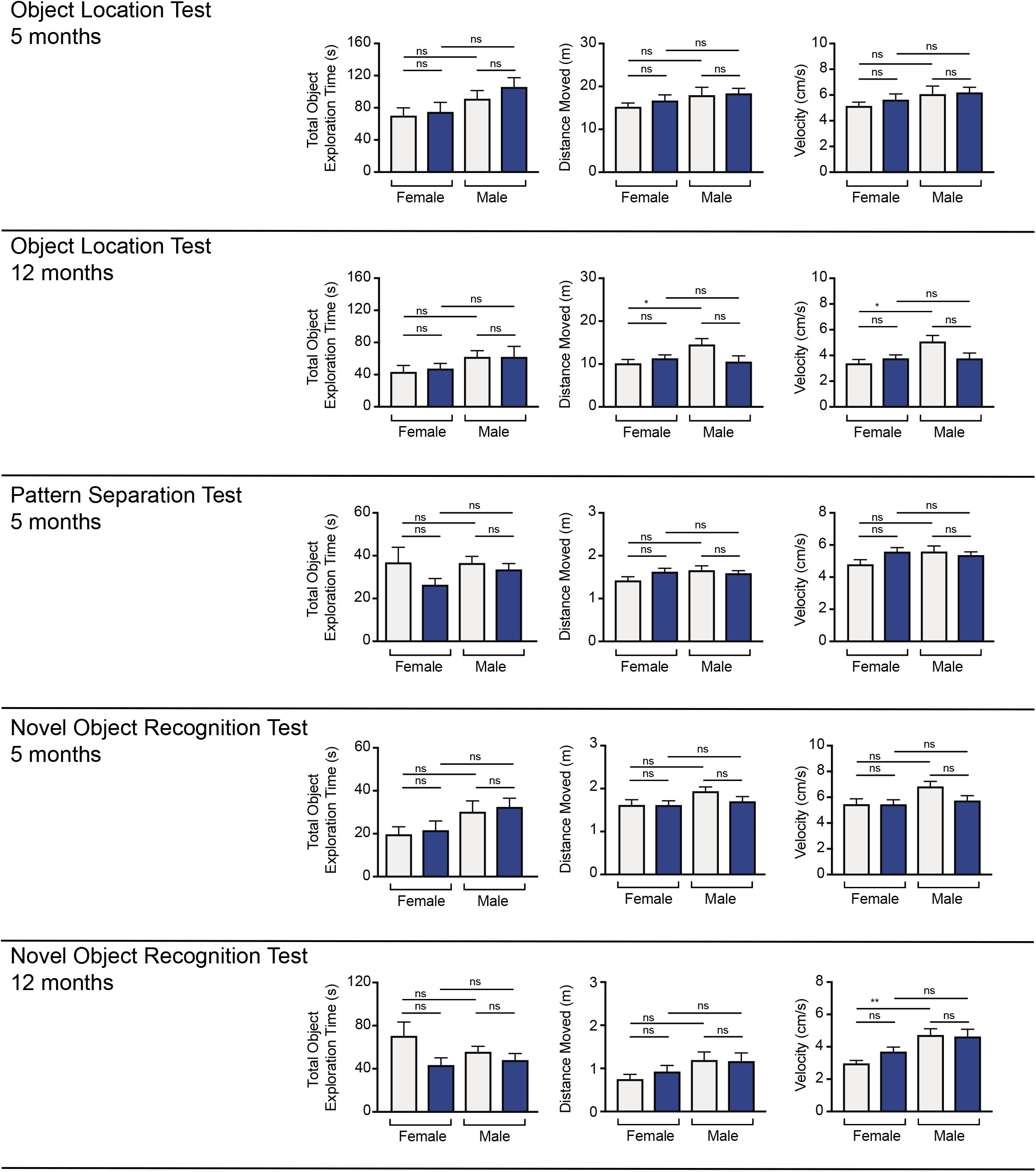
Control data for OLT (object location test), PST (pattern separation test), and NORT (novel object recognition test) experiments depicted in Figure 1: total object exploration time, distance moved, and velocity. For all the neurobehavioral tests (OLT, PST, and NORT) depicted in Figure1, data were collected on total object exploration time (seconds), distance moved (meters), and velocity (centimeters per second) of all mice for all three tests (object location test, OLT; pattern separation test, PST; novel object recognition test, NORT). Error bars represent the mean of biological replicates +/- SEM (n=9-10). Indicated p-values were calculated using one-way ANOVA with Tukey’s multiple comparison. *p < 0.05; **p < 0.01; ns, not significant.

**Supplemental Figure 2.**
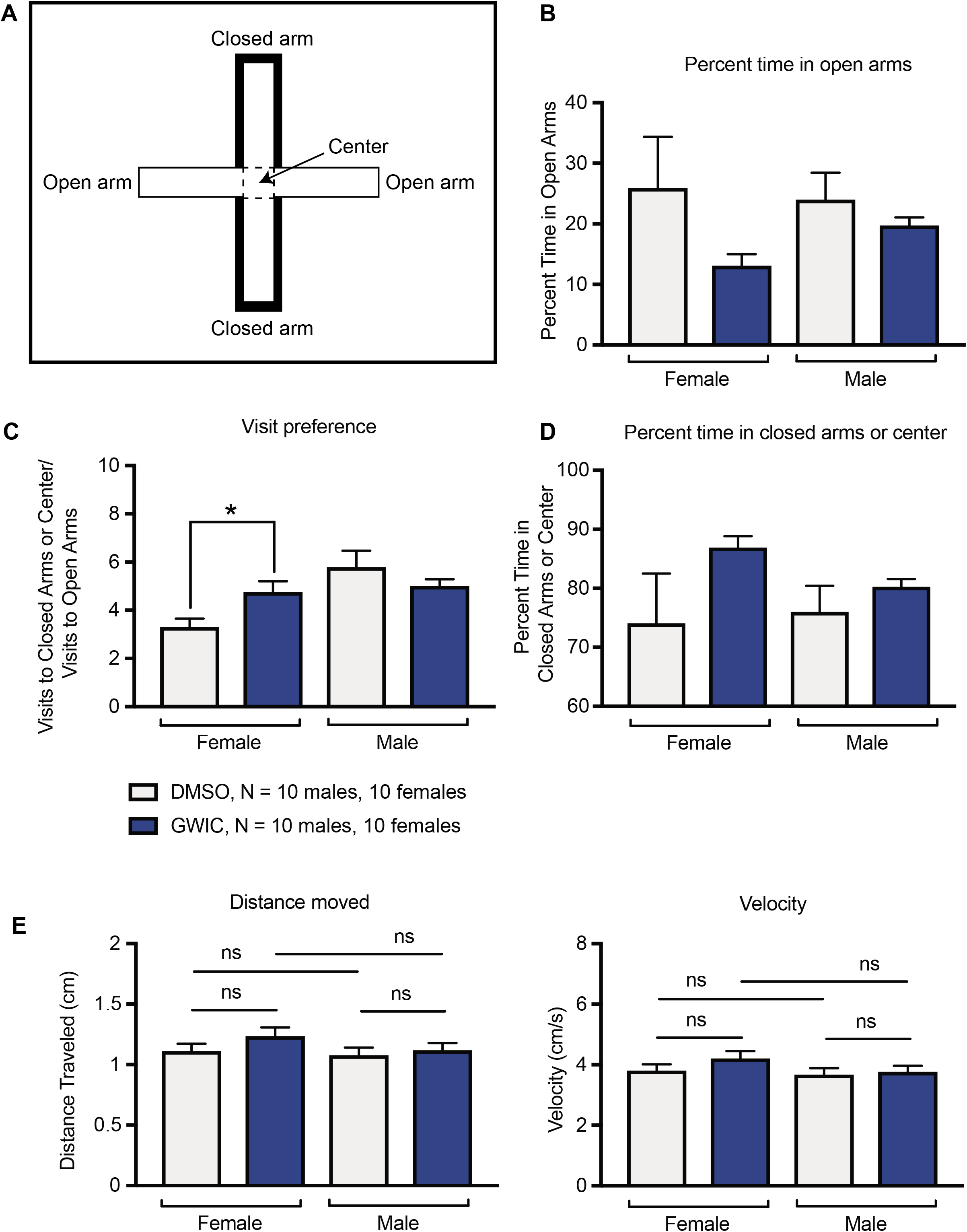
The elevated plus maze test (EPMT) was used at 5 months post-exposure to assess anxiety-related behavior in all cohorts. (A) The diagram depicts the experimental setup for this test: a 60 cm elevated plus-shaped apparatus with four arms, each 35 cm in length and 5 cm in width. Two of the arms are surrounded with walls on three sides of 15 cm in height (the two regions identified as “closed arm” in the diagram) and two of the arms are open (the two regions identified as “open arm” in the diagram). Mice are gently placed in the region identified as “center” in the diagram, facing a closed arm, and allowed to freely explore the maze for one five minute interval. (B) Percent time spent in the open arms was calculated and compared among cohorts. (C) Visits to each region were also calculated, as was visit preference, calculated as the ratio of number of visits to the closed arms or the center and the number of visits to the open arms. (D) Percent time spent in the closed arms or center was calculated and compared among cohorts. (E) Distance moved and velocity of all mice during the test. Error bars represent the mean of biological replicates +/- SEM (n=10). Indicated p-values were calculated using two-tailed, unpaired, Student’s t-test. *p < 0.05; ns, not significant.

**Supplementary Figure 3.**
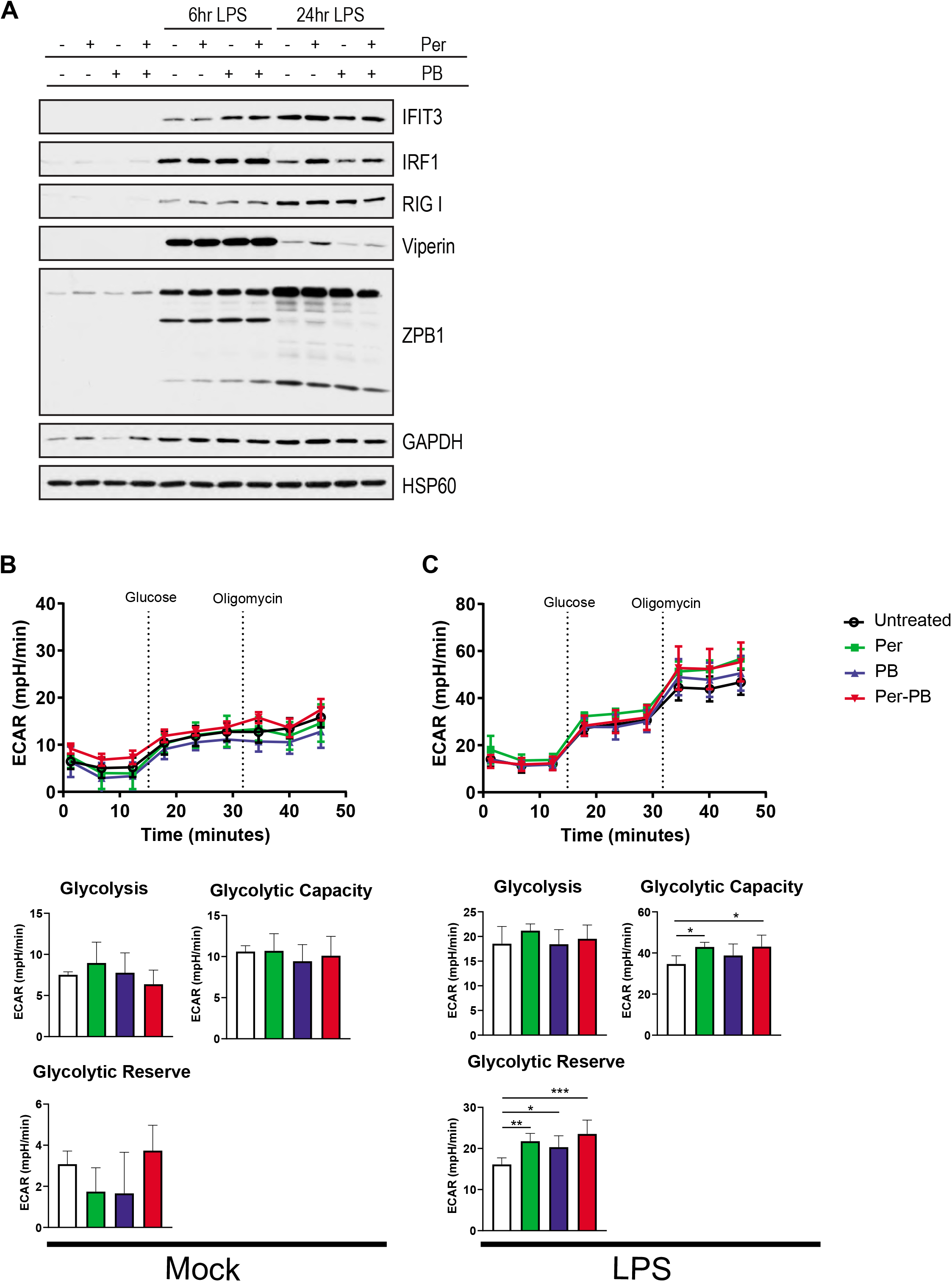
ISG expression, inflammasome activation, and glycolysis in BMDMs treated with GWIC. (A) Western blot analysis of ISG expression in BMDMs treated with overnight with GWIC, then treated with 200 ng/mL LPS for 6 or 24 hours as indicated. Cells extracts were blotted for indicated proteins as in Figure 2. (B) and (C) Glycolysis as measured by extracellular acidification rate (ECAR) in BMDMs treated with GWIC. BMDMs were treated overnight with GWIC as indicated. Cells were treated with Seahorse medium (Mock) (B) or 200 ng/mL LPS (LPS) for 6 hours (C), then ECAR was measured on a Seahorse XF96 Analyzer using a modified protocol as described in Section 2.8 of materials and methods. Error bars represent the mean of technical replicates +/- SD (n=5). Significance values for pairwise comparisons were calculated using the two-stage step-up method to yield q values that have been corrected for the false discovery rate. * q < 0.05; ** q < 0.01; *** q < 0.001; **** q < 0.0001.

**Supplemental Figure 4.**
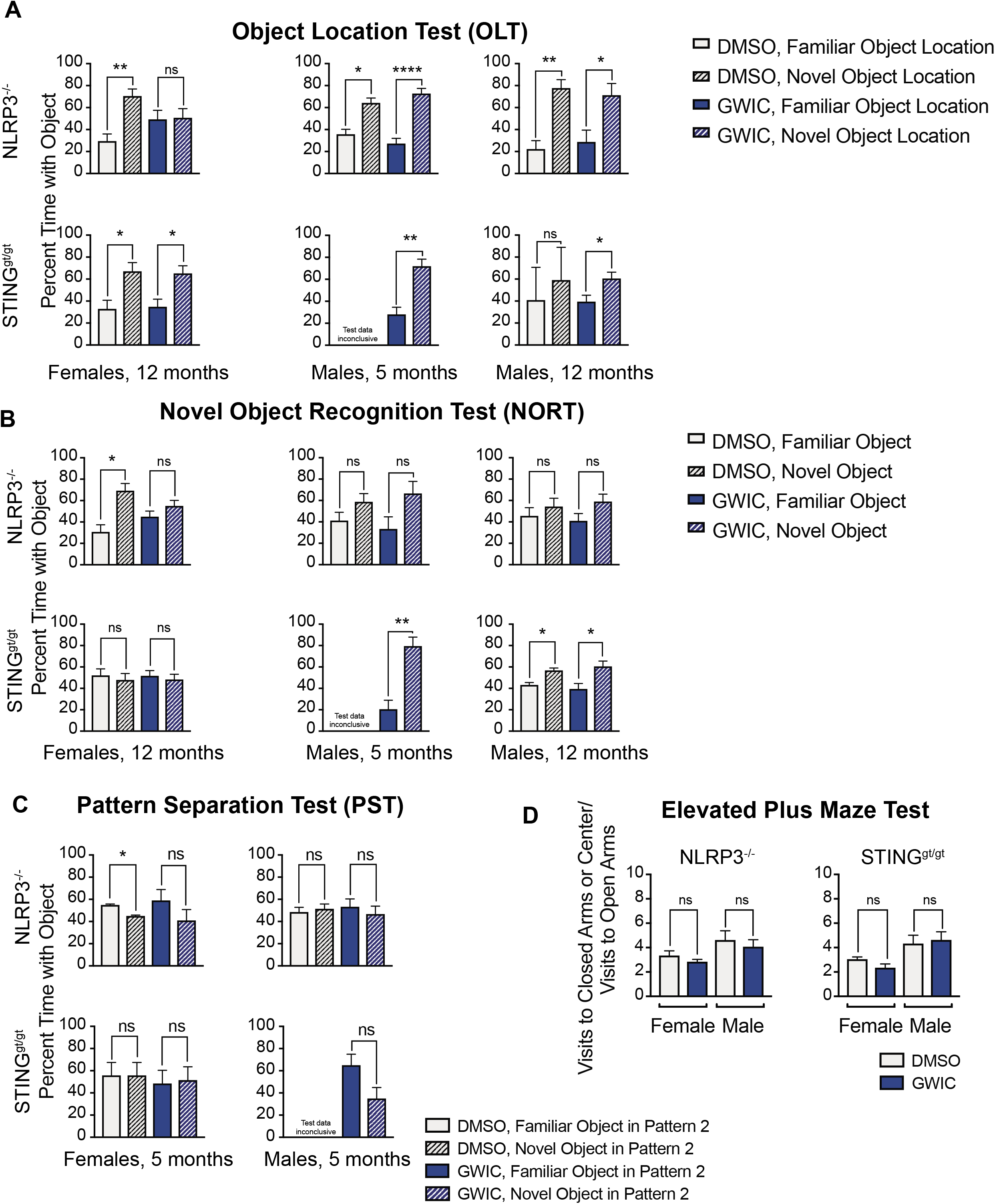
Addition neurobehavioral tests performed on DMSO- and GWIC-exposed NLRP3^−/−^STING^gt/gt^ C57BL/6 female mice. (A) Results of OLT testing on DMSO- and GWIC-exposed female NLRP3^−/−^and STING^gt/gt^ mice at 12 months post-exposure and DMSO- and GWIC-exposed male NLRP3^−/−^and STING^gt/gt^ mice at 5 and 12 months post-exposure. Experiments were performed as in Figure 1. (B) Results of NORT testing on DMSO- and GWIC-exposed female NLRP3^−/−^and STING^gt/gt^ mice at 12 months post-exposure and DMSO- and GWIC-exposed male NLRP3^−/−^and STING^gt/gt^ mice at 5 and 12 months post-exposure. Experiments were performed as in Figure 1. (C) Results of PST testing on DMSO- and GWIC-exposed female and male NLRP3^−/−^and STING^gt/gt^ mice at 5 and 12 months post-exposure. Experiments were performed as in Figure 1. (D) Results of EPMT testing on DMSO- and GWIC-exposed female and male NLRP3^−/−^and STING^gt/gt^ mice at 5 months post-exposure. Experiments were performed as in Supplemental Figure 2. Error bars represent the mean of biological replicates +/- SEM (n=5-10 for STING^gt/gt^ females and 3-6 for STING^gt/gt^ males, n=5-10 for NLRP3^−/−^females and 3-7 for NLRP3^−/−^males). Indicated p-values were calculated using two-tailed, unpaired, Student’s t-test. * p < 0.05; ** p < 0.01; **** p < 0.0001, ns, not significant.

**Figure S5.**
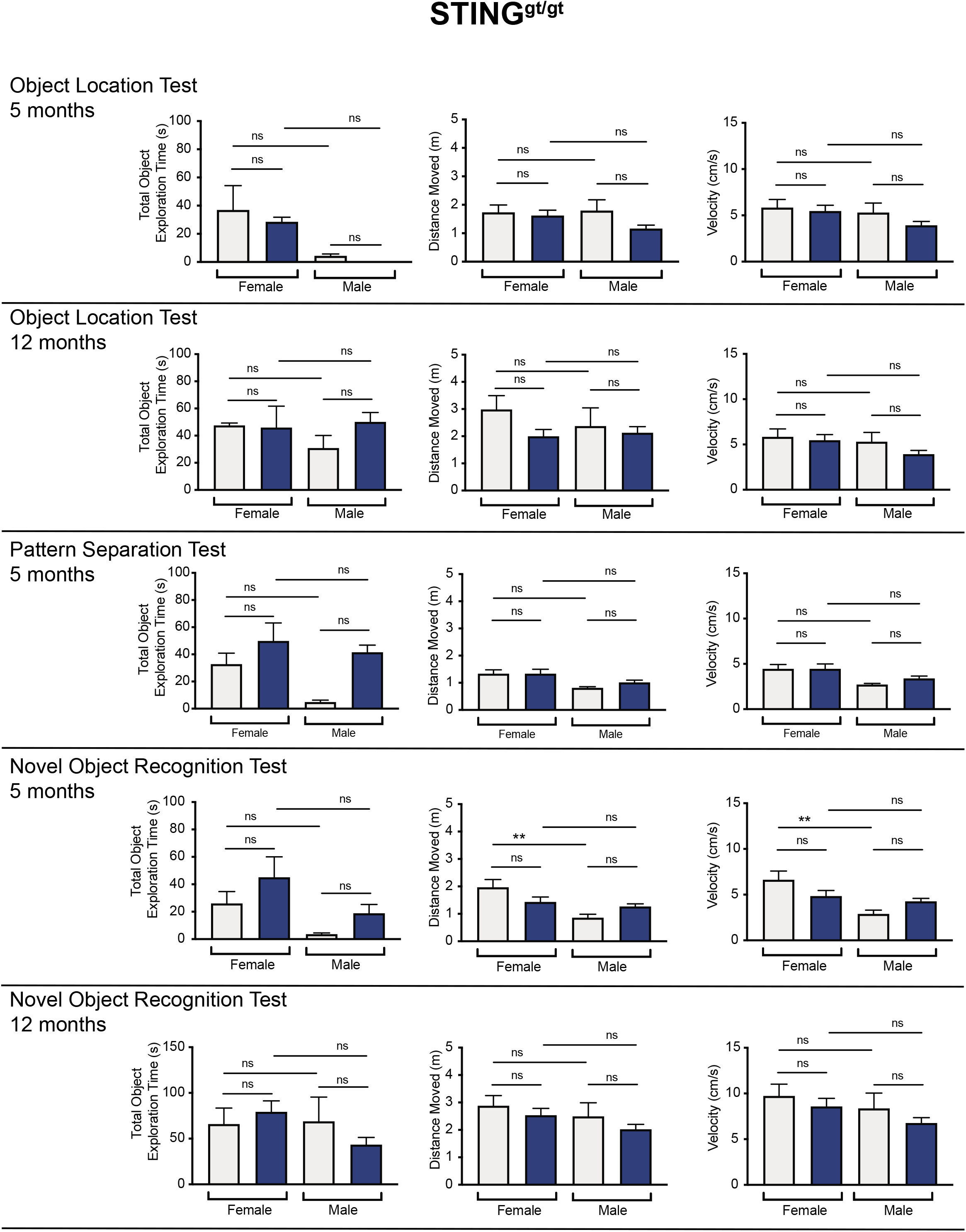
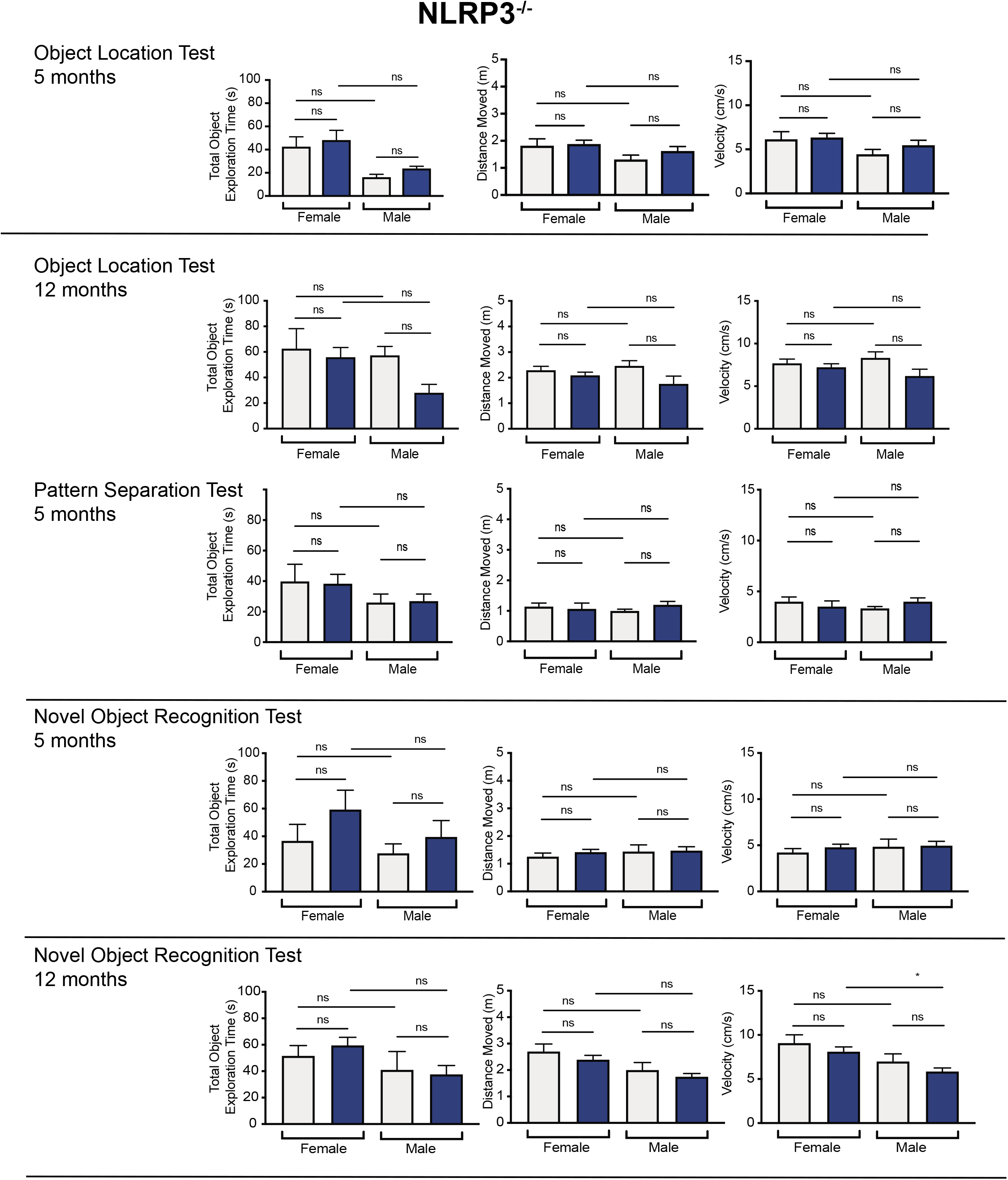
Control data for OLT (object location test), PST (pattern separation test), and NORT (novel object recognition test) experiments depicted in Figures 5 and 6: total object exploration time, distance moved, and velocity for STING^gt/gt^ and NLRP3^−/−^ mice. For the OLT, PST, and NORT data depicted in Figures 5, 6, and Supplemental Figure 4, data were collected on total object exploration time (seconds), distance moved (meters), and velocity (centimeters per second) of all mice for all three tests (object location test, OLT; pattern separation test, PST; novel object recognition test, NORT). Error bars represent the mean of biological replicates +/- SEM (n=5-10 for STING^gt/gt^ females and 3-6 for STING^gt/gt^ males, n=5-10 for NLRP3^−/−^females and 3-7 for NLRP3^−/−^males). Indicated p-values were calculated using one-way ANOVA with Tukey’s multiple comparisons. *p < 0.05; **p < 0.01; ns, not significant.

**Supplementary Figure 6.**
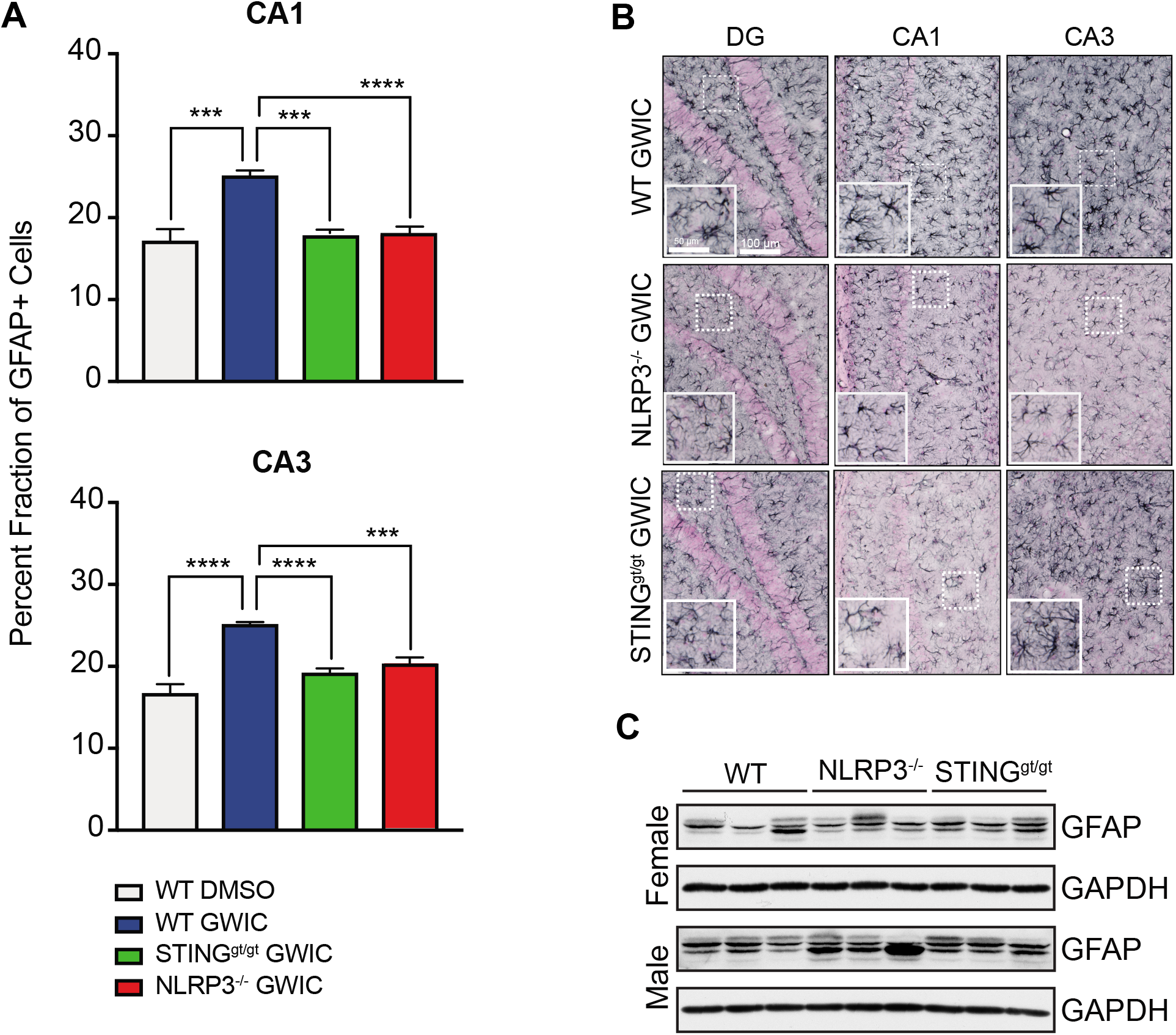
Percent fraction of GFAP+ cells and GFAP expression in the hippocampi of NLRP3^−/−^ and STING^gt/gt^ GWIC-exposed female mice. **(A)** Quantification of percent fraction GFAP+ cells within two regions of the hippocampus (the CA1 and CA3) in DMSO-exposed WT female mice and GWIC-exposed WT, NLRP3^−/−^ and STING^gt/gt^ female mice. Error bars represent the mean of biological replicates +/- SEM (n=6). Indicated p-values were calculated using two-tailed, unpaired, Student’s t-test. ***p < 0.001; **** p < 0.0001. (B) Images show representative distribution and morphology of GFAP+ astrocytes in these three regions in female GWIC-exposed WT, NLRP3^−/−^ and STING^gt/gt^ mice (scale bar = 100 μM). Insets show magnified views (scale bar = 50 μM). (C) Tissue extracts were generated from microdissected hippocampi from DMSO-exposed WT, NLRP3^−/−^ and STING^gt/gt^ mice and immunoblotted for GFAP as described in Materials and Methods. Each lane represents one mouse, with n=3 for each genotype and sex.

**Supplemental Figure 7.**
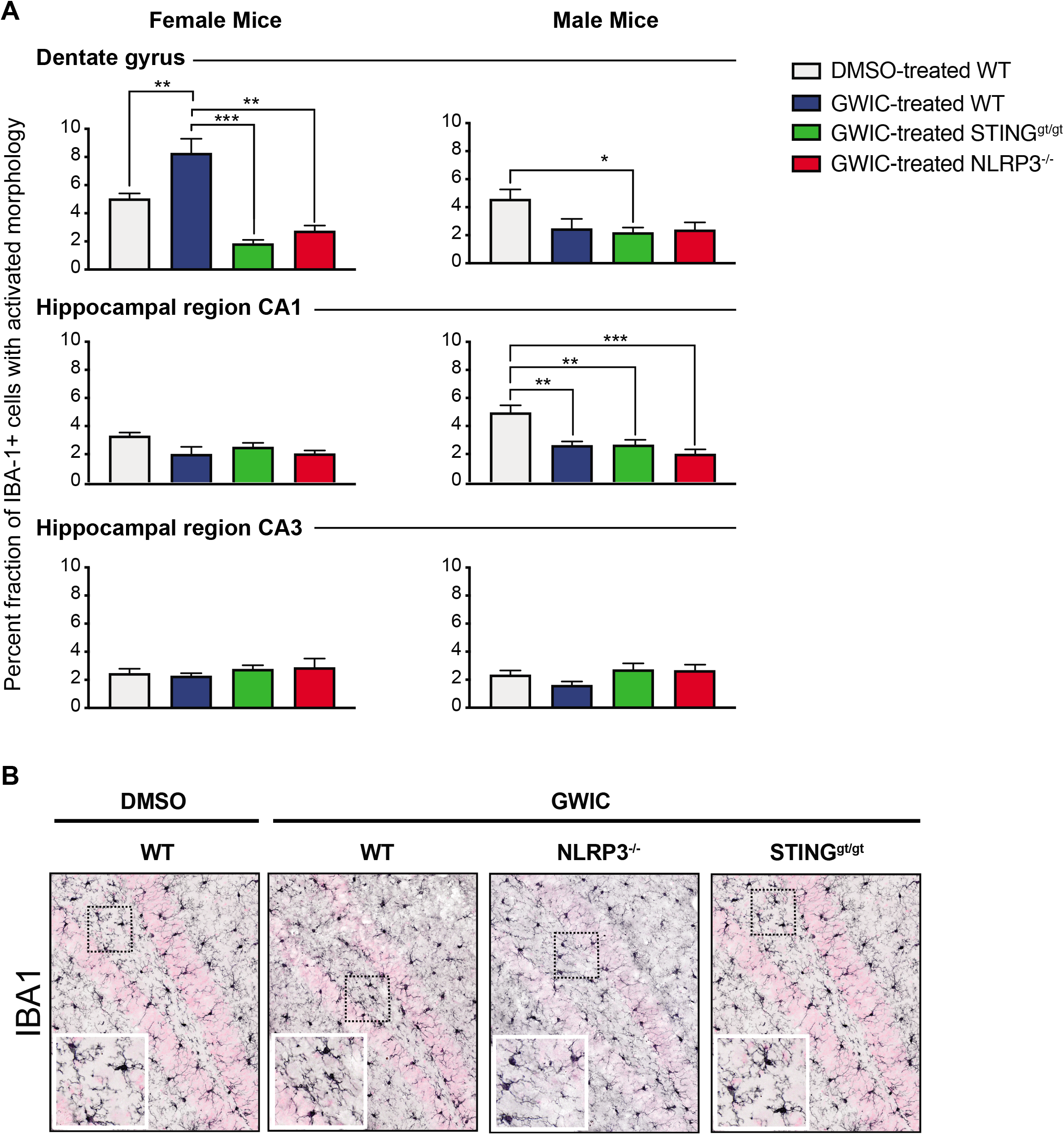
Analysis and imaging of IBA-1+ cells in the hippocampi of WT, NLRP3^−/−^ and STINGS^gt/gt^ mice. **(A)** Quantification of percent IBA-1+ cells with activated morphology in three hippocampal regions (DG, CA1, and CA3) in DMSO- and GWIC-exposed WT and GWIC-exposed NLRP3^−/−^ and STING^gt/gt^ female and male mice. Sections were scored blindly for total number of IBA-1+ cells and number of IBA-1+ cells with activated morphology. Error bars represent the mean of biological replicates +/- SEM (n=6). Indicated p-values were calculated using two-tailed, unpaired, Student’s t-test. *p < 0.05; **p < 0.01; **** p < 0.0001. Absence of statistical information indicates a lack of statistical significance. (B) Images show representative distribution and morphology of IBA-1+ microglia in the DG region of female DMSO- and GWIC-exposed WT, and GWIC-exposed NLRP3^−/−^ and STING^gt/gt^ mice (scale bar = 100 μM). Insets show magnified views (scale bar = 50 μM).

